# Blocking Interleukin-1β transiently limits left ventricular dilation and reduces cardiac lymphangiogenesis during pressure-overload in mice

**DOI:** 10.1101/2023.04.01.535056

**Authors:** C. Heron, T. Lemarcis, O. Laguerre, M. Bundalo, C Valentin, M. Valet, A. Dumesnil, JB. Michel, P. Mulder, A. Zernecke, V. Tardif, E. Brakenhielm

## Abstract

Blocking pro-inflammatory pathways, e.g. the inflammasome or interleukin (IL)-1β, is a promising therapeutic approach in heart failure (HF). We hypothesized that IL-1β may regulate cardiac lymphangiogenesis in response to chronic pressure-overload, and hence could impact the resolution of myocardial edema and inflammation and the development of cardiac fibrosis and HF.

We investigated cardiac, lymphatic, and immune effects of anti-IL-1β treatment during HF development following pressure-overload induced by transaortic constriction (TAC) in BALB/c mice. We also examined the impact of IL-1β on macrophages and lymphatic endothelial cells *in vitro*, and assessed links between perivascular fibrosis and lymphatics in HF patients.

We found that early anti-IL-1β treatment transiently increased cardiac infiltration of CD206^+^ macrophages and delayed left ventricular (LV) dilation, which however did not suffice to prevent HF development at 8 weeks post-TAC. In contrast, late anti-IL-1β treatment did not alter LV dilation, but reduced cardiac lymphangiogenesis. This was linked to a cell non-autonomous role of IL-1β in promoting cardiac lymphangiogenesis through stimulation of macrophage production and maturation of VEGF-C. Surprisingly, despite reduced lymphatic density in late anti-IL-1β-treated mice, cardiac inflammation, interstitial fibrosis, and HF development were not aggravated. Further, we found that perivascular lymphatic density, unaltered by anti-IL-1β, was negatively associated with perivascular fibrosis in HF patients and our TAC model.

In conclusion, IL-1β blockage elicited transient functional cardiac benefit when initiated before LV dilation post-TAC in mice. In contrast, late treatment reduced cardiac lymphangiogenesis but did not impact HF development. Our study suggests that the therapeutic window for anti-IL-1β treatment may be crucial, as initiation of treatment during the late lymphangiogenic response, induced by LV dilation, may diminish the potential cardiac benefit in HF patients. Finally, our data support a role of perivascular lymphangiogenesis in limiting perivascular fibrosis.

## Introduction

Interleukin-1β (IL-1β) signaling has emerged as a promising therapeutic target to limit inflammation in cardiovascular diseases, as highlighted in the CANTOS study, where *Canakinumab* reduced major adverse cardiovascular events in atherosclerotic patients.(*1*) Currently, other IL-1β blocking agents, including *Anakinra*, are being evaluated in the setting of ischemic heart failure (HF)(*2*). While cardiac IL-1β levels are increased also in non-ischemic HF, e.g. in dilated cardiomyopathy (DCM)(*3*), fewer experimental studies have investigated the impact of IL-1β-targeting in these settings. Thus, the potential of IL-1β as a therapeutic target to limit HF development in non- ischemic cardiomyopathy remains uncertain.

IL-1β not only regulates inflammation and fibrosis (in part by stimulating IL-6), but also impacts cardiac function *directly* due to negative inotropic effects(*4*). In addition, IL-1β may stimulate cardiomyocyte hypertrophy *via* upregulation of insulin-like growth factor (*Igf1*)(*5*). Most recently, CC chemokine receptor-2 (CCR2)-expressing myeloid cells have been suggested as a key source of IL-1β in failing human hearts, and found to be essential for driving cell-fate commitment of cardiac fibroblasts into periostin (Postn)-expressing matrifibroblasts mediating interstitial fibrosis after chronic pressure-overload by transverse aortic constriction (TAC) in mice(*6, 7*). Promisingly, reduction of cardiac IL-1β production during pressure-overload, by early treatment with a selective pharmacological inflammasome (NLRP3) inhibitor, reduced both cardiac hypertrophy and fibrosis at 5 weeks post-TAC in male C57Bl/6J mice.(*8*) However, another study, employing an IL-1β-blocking antibody post-TAC, demonstrated that the reduction in left ventricular (LV) hypertrophy led to accelerated LV dilation and HF development(*9*), potentially linked to reduced IL-1β-induced VEGF-A-mediated cardiac angiogenesis in response to cardiac hypertrophy. Similarly, a recent study, based on *Nlrp3* gene deletion in male C57Bl/6J mice, also reported that reduced inflammasome activation limited early compensatory cardiac hypertrophy following TAC, resulting in accelerated LV dilation and aggravated cardiac dysfunction(*10*).

Recently, we demonstrated that while pressure-overload leads to compensated concentric hypertrophy in female C57Bl/6J mice, in BALB/c mice, characterized by elevated cardiac *Il1b* gene expression post-TAC, it leads to eccentric hypertrophy with severe LV dilation.(*11*) Moreover, we demonstrated that LV dilation-induced wall stress, but not LV hypertrophy *per se*, triggers massive cardiac lymphangiogenesis, which however fails to limit cardiac inflammation and edema.(*11*) Lymphatics are essential for cardiac function, and consequently cardiac lymphatic dysfunction, or insufficient lymphangiogenesis, has been shown experimentally to contribute to adverse cardiac remodeling, inflammation, and fibrosis following myocardial infarction (MI) or pressure-overload.(*12–17*) Conversely, inflammatory mediators, including IL- 1β, may modulate lymphangiogenesis and/or lymphatic function.(*18–20*) Interestingly, we previously observed a striking correlation between cardiac *Il1b* and *Vegfc* expression(*11*), in line with previous reports indicating both direct and indirect pro-lymphangiogenic effects of IL-1β in other organs. This includes stimulation of VEGF- C-mediated proliferation of lymphatic endothelial cells (LEC) *via* an autocrine loop(*18*), as well as IL-1β-mediated activation of VEGF-C-release by macrophages to promote lymphangiogenesis during chronic inflammation(*21*). In contrast, several studies have demonstrated that local inflammation induces lymphatic dysfunction, as pro-inflammatory mediators, including IL-1β, act on lymphatic smooth muscle cells to reduce lymphatic drainage.(*22*) Additionally, in the heart, where lymphatic drainage depends directly on the force of cardiac contraction(*23*), the negative inotropic effects of IL-1β may indirectly reduce cardiac lymphatic transport capacity in cardiovascular diseases.

Here, we investigated the impact of anti-IL-1β treatment on cardiac lymphatics, inflammation, and LV remodeling during HF development post-TAC in BALB/c mice. We hypothesized that elevated cardiac *Il1b* may provide a molecular link between increased wall stress and cardiac lymphangiogenesis in response to pressure-overload. Indeed, cardiac wall stress (induced by LV dilation) likely triggers cardiac fibroblast IL-1β secretion, which is sensitive to mechanical stress(*5*). Specifically, we argued that the striking lymphangiogenesis observed in our TAC model may be indirectly induced by elevated cardiac IL-1β driving local VEGF-C production by cardiac cells. Thus, we expected that interfering with IL-1β signaling post-TAC would reduce cardiac lymphangiogenesis, leading to aggravation of inflammation and fibrosis and accelerated HF development. To investigate the cardiac functional and lymphatic impact of IL-1β in different stages of pressure-overload-induced HF, we applied an anti-IL-1β antibody (*Gevokizumab*) before and after development of LV dilation induced by TAC in BALB/c mice (*early* and *late* treatment, **Fig. 1a**). To dissect the cellular mechanisms of IL-1β in regulation of lymphangiogenesis, we investigated its molecular impact *in vitro* in macrophage and LEC cultures. Finally, we investigated whether lymphatics in the perivascular niche may be linked to perivascular fibrosis in mice and humans.

**Figure 1.**
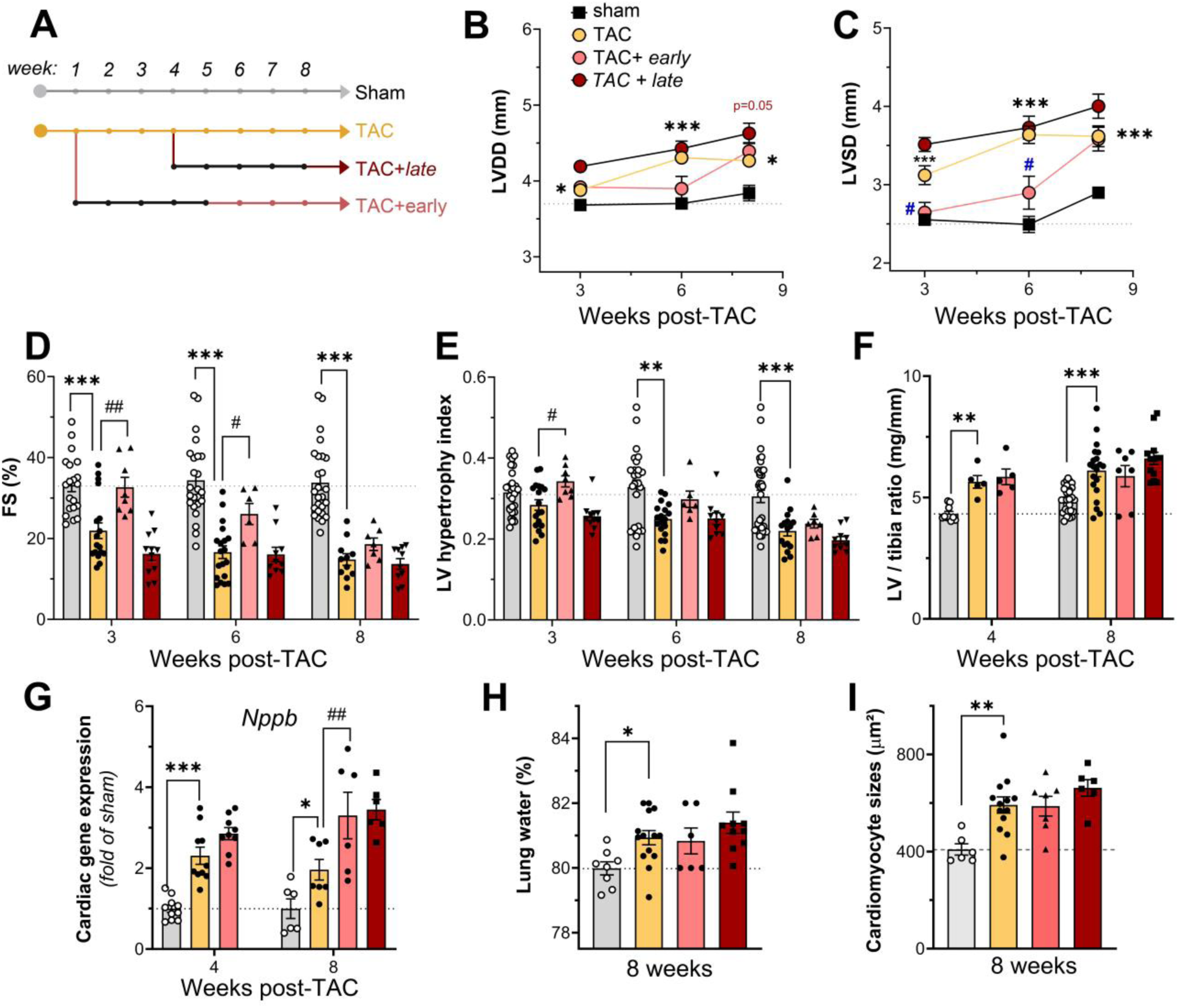
Early anti-IL-1β treatment transiently delays LV dilation but does not impact HF development by 8 weeks post-TAC. Schematic overview of the study (**a**). Echocardiography was used at 3, 6, and 8 weeks post-TAC to assess LV diastolic diameter (*LVDD*, **b**) and LV systolic diameter (*LVSD*, **c**), and to calculate fractional shortening (*FS*, **d**) in sham (*n=29;* grey bars w. white circles), control TAC (*n=18;* yellow bars w. black circles), and *early* (*n=8;* red bars w. black circles) vs. *late* (n*=10;* red bars w. black squares) anti-IL-1β-treated TAC mice. LV hypertrophy dilatation index (**e**) Left ventricular (LV) weights normalized to tibia lengths at 4 and 8 weeks (**f**). Cardiac expression analyses of *Nppb* at 4 and 8 weeks (**g**, *n= 6-10* per group). Lung water content at 8 weeks (**h**). Cardiomyocyte cross-sectional area at 8 weeks (**i**, *n=6-13* animals per group). Mean ± s.e.m are shown. Groups were compared by two-way ANOVA followed by Dunnett’s multiple comparison test (panels **b-g**), by one-way ANOVA followed by Sidak’s multiple comparison test (**h**), or by non-parametric Kruskal Wallis followed by Dunn’s posthoc test (**i**) * p<0.05, ** p<0.01, *** p<0.001 *versus* sham; # p<0.05; ## p<0.01 *versus* TAC control.

## Material and Methods

### Study approval

Animal housing and experiments were in accordance with European Directive 2010/63/EU on the protection of animals, and the study was approved by the Normandy University ethical review board Cenomexa according to French and EU legislation (APAFIS #**23175**-2019112214599474 v6; APAFIS #**32433**-2022070712508369 v2). A total of 117 BALB/c female mice, surviving TAC or sham-surgery were included in this study. Anonymized human heart samples evaluated in this study were obtained from the Bichat hospital biobank (*U1148 BB-0033-00029/* BBMRI, coordinator JB Michel) authorized for tissue collection by the Inserm institutional ethical review board, as previously reported(*11*). All samples were collected in accordance with national legislation and applicable codes of conduct: Charter of the Fundamental Rights of the EU; the Declaration of Helsinki (2008).

### Experimental mouse model, macrophage cell culture, and human samples

TAC surgery was performed in adult female BALB/c mice (Janvier Laboratories, France). Mice were anesthetized by intraperitoneal injection of ketamine (100 mg/kg) and xylazine (10 mg/kg). Briefly, a minimally invasive method was used to constrict the aortic arch, using a 26G needle. Double-banding of the aorta was applied to prevent suture internalization and model variability, as described.(*11*) Buprenorphine (50 µg/kg) was injected subcutaneously 6 hours after surgery and twice per day until 3 days post-operation. Anti-IL-1β treatment consisted of repeated intraperitoneal injections of a blocking monoclonal antibody against IL-1β, (Gevokizumab/XOMA-052 provided by Servier, France), as previously described(*24*), using a dose of 200 µg/mouse (10 mg/kg) three times per week, starting from 7 days post-TAC (*week 1*) until week-5 post-TAC (*early* treatment), or beginning in week-4 until week-8 post-TAC (*late* treatment). Euthanasia was performed by pentobarbital overdose (100 mg/kg Euthoxin). For details see *Suppl. Methods*.

Bone marrow-derived macrophages (BMM) were prepared from C57BL/6 mice and cultured, as described(*25*). Briefly, serum-starved BMMs were seeded in 6-well plates (6×10^6^/well) and exposed to 10 ng/mL recombinant mouse IL-1β (PeproTech). After 4h or 24h of culture, cells were recovered and RNA extracted for qPCR analyses. Balb/c dermal LECs (CellBiologics, BALB-5064L) were seeded in 6-well plates (3×10^5^/well) and grown to confluence, serum-starved overnight, and exposed to 20 ng/mL recombinant mouse IL-1β (PeproTech). After 24h, cells were recovered and RNA extracted for RNAseq (Novogene). For details see *Suppl. Methods*.

Cardiac samples from end-stage HF patients (recipients of cardiac transplants at the Bichat Hospital in Paris, France) were examined by histology (Sirius Red staining). Discarded cardiac autopsy samples, obtained from age-matched donors without cardiovascular disease, were used as healthy controls, as previously reported(*11*). This study was limited to retrospective analyses of samples from a preexisting biobank (*U1148 BB-0033-00029/* BBMRI) and did not include collection of new samples from patients.

### Cardiac functional, cellular, and molecular analyses

Cardiac function was evaluated by echocardiography as described.(*11*) Cardiac sections were analyzed by histology and immunohistochemistry, as described,(*11*) to determine lymphatic vessel densities and sizes, immune cell infiltration (macrophages and T cells), cardiomyocyte hypertrophy, and fibrosis (Sirius Red staining). Perivascular fibrosis was defined as the area of fibrosis surrounding the media in similar-sized arterioles, as described.(*26*) Cardiac gene expression was analyzed by RT-qPCR. Cardiac protein levels of VEGF-C and VEGF-D were assessed by Western Blot. Cardiac and plasma levels of IL-1β were evaluated by ELISA (RnD Systems: ELISA DuoSet anti-IL-1β /IL-1F2 #DY401-05). For details see *Suppl. Methods*.

### Statistics

Data are presented as mean ± s.e.m. Comparisons were selected to determine: 1) impact of pathology (healthy sham *vs*. TAC); 2) effect of treatment (anti-IL-1β-treated vs. TAC control); and 3) effect of IL-1β vs. untreated control cells. Statistical analyses for comparisons of two independent groups were performed using either Student’s two-tailed t-test for groups with normal distribution, or alternatively by Mann Whitney U test for samples where normality could not be ascertained based on D’Agostino & Pearson omnibus normality test. For comparisons of three groups or more either one-way ANOVA followed by Sidak’s multiple comparison posthoc (for parameters with n>7 with normal distribution), or alternatively Kruskal-Wallis nonparametric analysis followed by Dunn’s posthoc for multiple comparison (for parameters with non-Gaussian distribution) were performed. Longitudinal echocardiography studies were analyzed by paired two-way ANOVA followed by Dunnett’s posthoc, while morphometric data and gene expression data were analyzed by two-way ANOVA followed by Sidak’s posthoc for multiple pair-wise comparisons or Dunnett’s posthoc to compare 3 or more groups. Non-parametric Spearman rank order test was used to evaluate correlations. Outlier samples were identified as an individual value exceeding group mean±4SD in groups with n<u>≥</u>7. All statistical analyses were performed using GraphPad Prism software.

## Results

### 1. Early, but not late, anti-IL-1β treatment delays LV dilation and dysfunction but does not limit HF development post-TAC

Evaluation of cardiac function by echocardiography at 3-, 6-, and 8-weeks post-TAC demonstrated that *early* anti-IL-1β treatment transiently prevented LV dilation and dysfunction up to 6 weeks-post-TAC (**Fig. 1b-e**; **Table 1**; **Suppl. Table S1**). Indeed, the reduction in LV hypertrophy index, observed in TAC controls starting from 3 weeks, was delayed in *early*-treated mice (**Fig. 1e**). However, cardiac hypertrophy was not significantly altered as compared to TAC controls (**Fig. 1f**; **Suppl. Fig. 1**), although cardiac gene expression of *Nppb* (encoding BNP) was elevated at 8 weeks in *early*- treated mice as compared to TAC controls (**Fig. 1g**). The transient beneficial effect on cardiac function of antibody-mediated IL-1β blockage was lost at 8 weeks, when *early*- treated animals exhibited a similar reduction in fractional shortening (FS) as TAC controls (**Fig. 1d**). Gravimetric measurements at 8 weeks post-TAC revealed no effects of *early*-stage treatment on development of pulmonary edema (**Fig. 1h**).

**Table 1.**
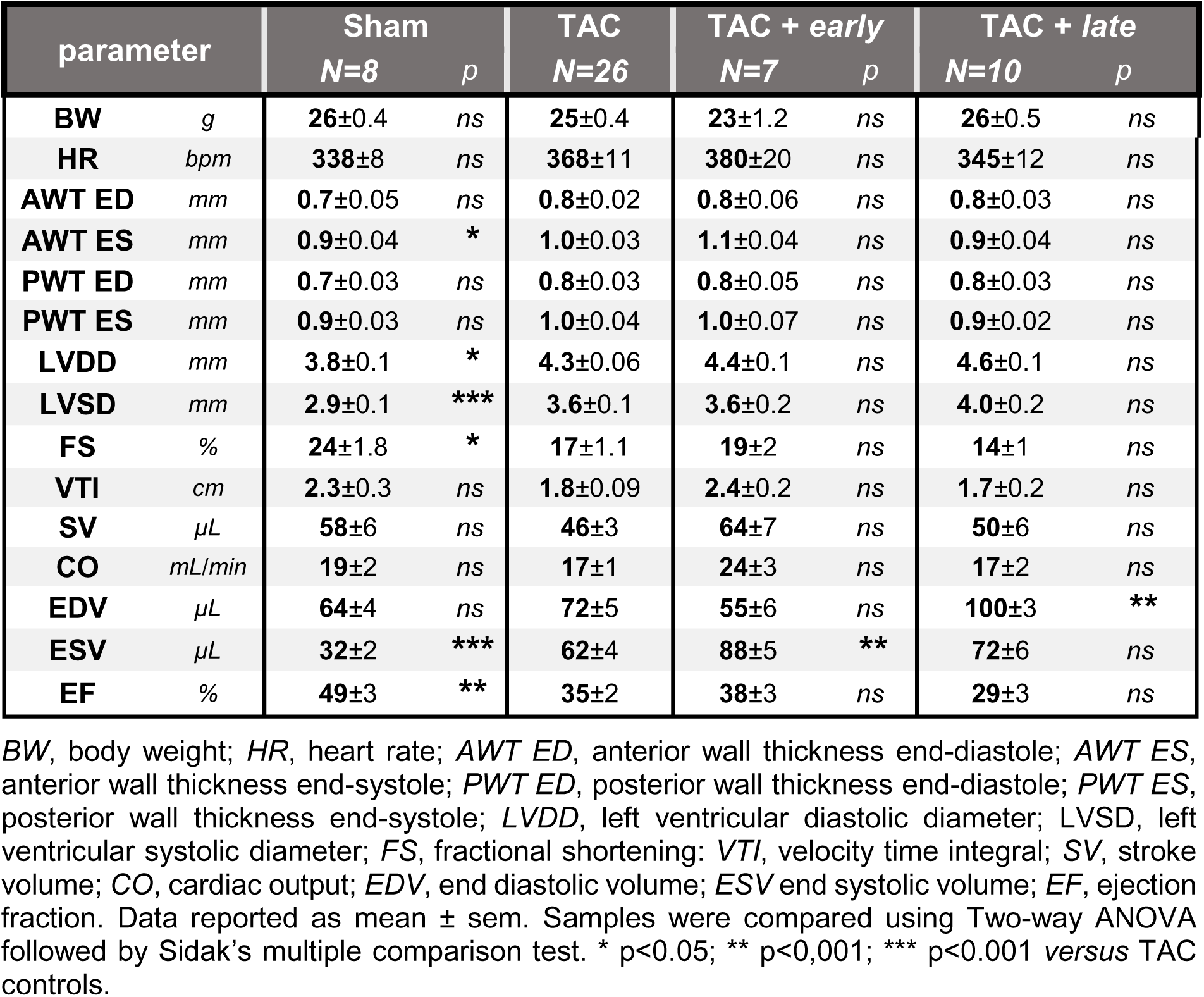
Echocardiography at 8 weeks post-TAC in BALB/c mice.

In contrast, *late* anti-IL-1β treatment did not alter LV dilation (**Fig. 1b, c**, **Table 1**), cardiac dysfunction, hypertrophy, or pulmonary edema at 8 weeks (**Fig. 1d-h**). Evaluation of cardiomyocyte sizes revealed moderate cardiac hypertrophy at 8 weeks post-TAC, which was unaltered by anti-IL-1β treatment (**Fig. 1i**).

### 2. E*arly* anti-IL-1β treatment moderately affects cardiac inflammation post-TAC

We and others have previously shown biphasic upregulation of cardiac IL-1β during experimental pressure-overload, with a rapid increase (from 3 days post-TAC^21^), followed by a second progressive increase (from 3-8 weeks post-TAC).(*11, 27*) Our cardiac immunohistochemical analyses revealed increased cardiac IL-1β levels, notably in areas of macrophage accumulation and interstitial fibrosis (**Suppl. Fig. 2a**). Pressure-overload not only activates inflammatory pathways in the heart, but also systemic innate immune cell activation, which in the chronic setting leads to splenomegaly. (*27*),(*28*) We found that both *early* and *late* anti-IL-1β treatment significantly reduced spleen weight at 8 weeks post-TAC (**Fig. 2a**), reflecting systemic anti-inflammatory effects of the treatment, as we previously described in rats.(*24*) Only *late* anti-IL-1β treatment led to a compensatory increase in circulating IL-1β plasma levels, as evaluated by ELISA (**Fig. 2b**). Of note, rather than neutralizing the cytokine, *Gevokizumab* acts by reducing IL-1β binding to the IL-1R receptor.(*24*) Neither *early* nor *late* treatment altered cardiac IL-1β gene or protein levels post-TAC (**Fig. 2c, d**; **Suppl. Fig. 2a**). Unexpectedly, anti-IL-1β treatment did not reduce cardiac gene expression of other proinflammatory cytokines, such as IL-6, as evaluated by qPCR (**Fig. 2e**), and both *Il1b* and *Il6* cardiac expression levels correlated with the degree of LV dilation at 8 weeks post-TAC (**Suppl. Fig. 2b, c**). Intriguingly, *early* anti-IL-1β treated mice displayed increased cardiac levels of both *Ccl2* and *Ccl7* at 4 weeks as compared to control TAC mice (**Fig. 3a**).

**Figure 2.**
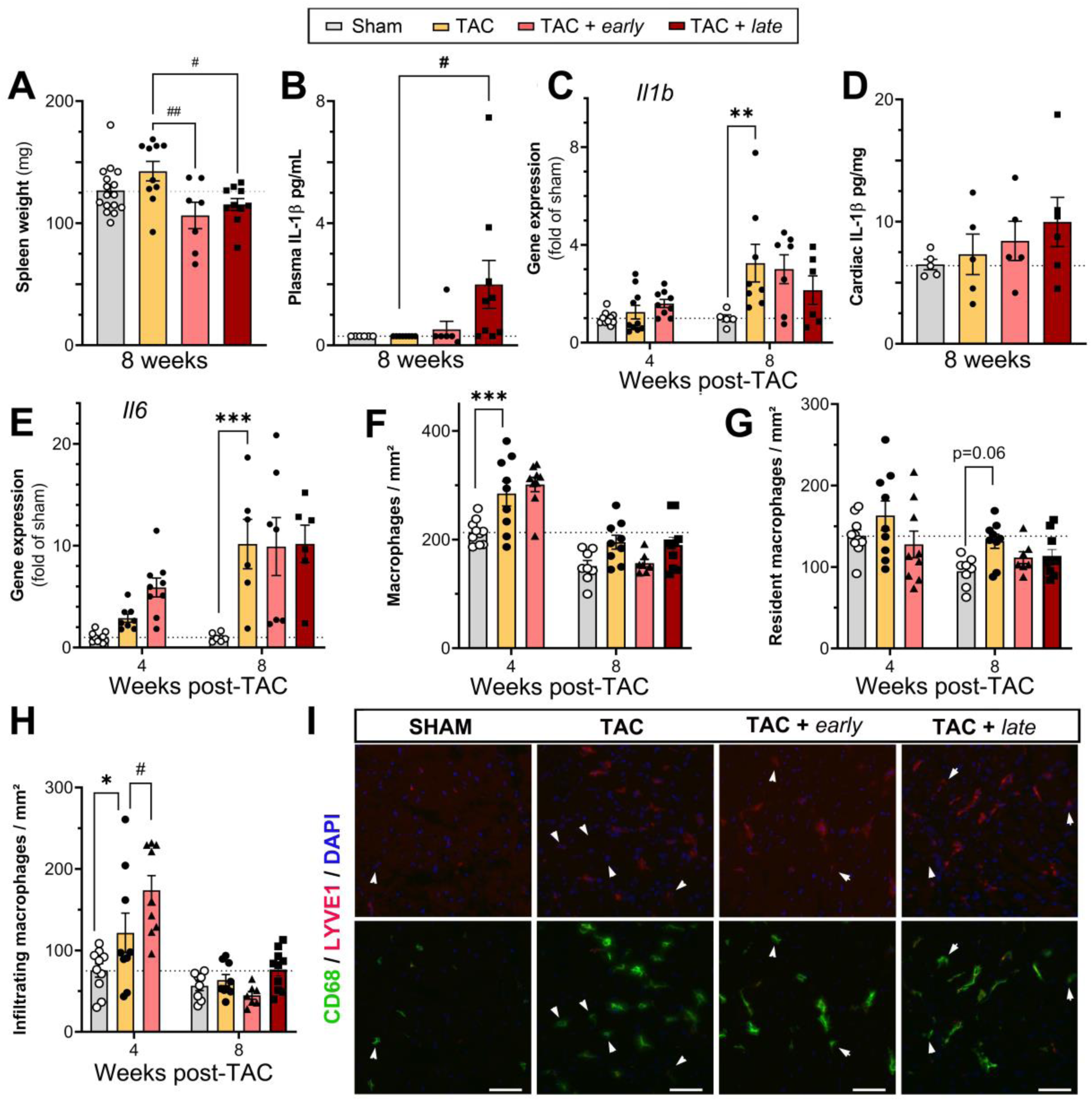
Early anti-IL-1β treatment does not reduce cardiac inflammation. Spleen weights at 8 weeks (**a**) in sham (*n=16;* grey bars w. white circles), control TAC (*n=10;* yellow bars w. black circles), and *early* (*n=7;* red bars w. black circles) vs. *late* (n*=10;* red bars w. black squares) anti-IL-1β-treated TAC mice. Plasma IL-1β (**b**) evaluated by ELISA at 8 weeks (*n=5-10* mice per group). Cardiac *Il1b* expression (**c**) evaluated by qPCR at 4 and 8 weeks, and cardiac IL-1β protein levels (**d**) evaluated by ELISA at 8 weeks. Cardiac *Il6* expression (**e**) evaluated by qPCR at 4 and 8 weeks (*n=5-10* mice per group). Total cardiac CD68^+^ macrophage levels (**f**); tissue-resident CD68^+^ / Lyve1^+^ macrophages (**g**); and infiltrating CD68^+^Lyve1^neg^ macrophage subpopulations (**h**) evaluated by immunohistochemistry at 4 and 8 weeks (*n=7-10* mice per group). Examples of cardiac macrophages (**i**). CD68, *green*, Lyve1, *red*, Dapi *blue*, Scalebar 50 µm. Arrowheads point to infiltrating (non-resident) CD68^+^/Lyve1^neg^ macrophages. Mean ± s.e.m are shown. Groups were compared by one-way ANOVA followed by Sidak’s multiple comparison test (**a, d**), by non-parametric Kruskal Wallis followed by Dunn’s multiple comparison posthoc test (**b**), or by two-way ANOVA mixed-effect analysis followed by Dunnett’s multiple comparison test (**c, e-h**). * p<0.05, *** p<0.001 *versus* sham; # p<0.05; ## p<0.01 *versus* TAC control.

**Figure 3.**
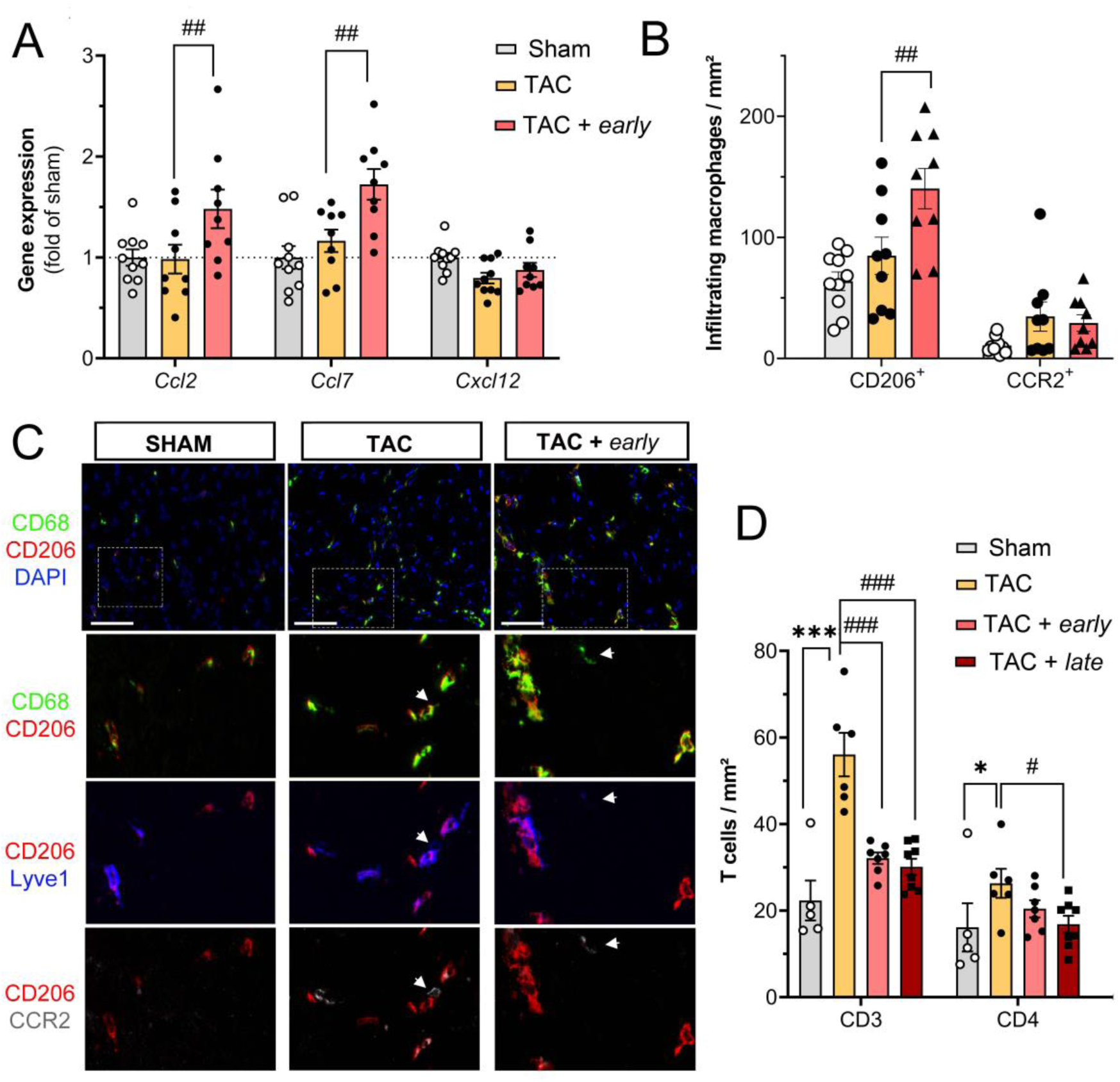
Differential effects of anti-IL-1β treatments on cardiac recruitment of macrophage subpopulations and T cells. Cardiac expression analyses (qPCR) of chemokines at 4 weeks in sham (*n=10;* grey bars w. white circles), control TAC (*n=5;* yellow bars w. black circles), and *early* (*n=9;* red bars w. black circles) vs. *late* (n*=9;* red bars w. black squares) anti-IL-1β-treated TAC mice (**a**). Quantification at 4 weeks post-TAC of cardiac-infiltrating (Lyve1^neg^) macrophage subpopulations positive for either CD206 or CCR2 (**b**). Examples of cardiac macrophages (**c**). *1^st^ row*: CD68 (*green*) CD206 (*red*) DAPI (*blue*). Scalebar 50 µm. (The white hashed areas are shown magnified in rows 2-4 of panel b). Resident macrophages are LYVE1^+^ (*blue, 3^rd^ row*), and most are CD206^+^. Cardiac-infiltrating (non-resident) macrophages are negative for Lyve1 and positive for either CD206 (*red*) or CCR2 (*grey, white arrowhead*). Quantification of cardiac T cells at 8 weeks, including total CD3^+^ cells, and CD4^+^ subpopulations in sham (*n=3*), control TAC (*n=6*), and *early* (*n=7*) vs. *late* (n*=8*) anti-IL-1β-treated TAC mice. (**d**). Data reported as fold of control (*a*) or cells/mm² (*c, d*) (mean ± sem). Groups were compared by two-way ANOVA followed by Dunnett’s multiple comparison test, * p<0.05; *** p<0.001 *versus* sham. # p<0.05, ## p<0.01 *versus* TAC controls.

Concerning the impact on cardiac immune cells post-TAC, neither *early* nor *late* treatment significantly altered total (CD68^+^) or cardiac resident (CD68^+^/Lyve1^+^) macrophage levels at 4 or 8 weeks (**Fig. 2f, g, i**), as evaluated by immunohistochemistry. However, *early* anti-IL-1β treatment transiently increased levels of CD68^+^/Lyve-1^neg^/CD206^+^ macrophages (**Fig. 2h**), while not affecting CD68^+^/Lyve-1^neg^/CCR2^+^ macrophages (**Fig. 3b, c**) despite increased cardiac *Ccl2* expression.

Concerning adaptive immune cells, we previously showed, using flow cytometry, low cardiac B cell levels and elevated CD4^+^ T cells at 8 weeks post-TAC.(*11*) Examining cardiac T cells levels in anti-IL-1β treated groups, we found a decrease, rather than an increase, in total cardiac CD3^+^ T cells in both *early* and *late-*treated TAC mice (**Fig. 3d**).

### 3. Anti-IL-1β treatment alters cardiac lymphangiogenesis post-TAC

We previously demonstrated that cardiac *Vegfc* gene expression increased starting at 4 weeks post-TAC in BALB/c mice, leading to stimulation of cardiac lymphangiogenesis during the subsequent decompensation phase (5-8 weeks post-TAC), characterized by LV dilation.(*11*) We found that *early*-treatment, despite transiently increasing cardiac-infiltrating CD206^+^/Lyve1^neg^ macrophages, a potential source of VEGF-C post-TAC(*29*), did not alter cardiac *Vegfc* levels, nor lymphatic markers (*Lyve1, Flt4, Pdpn, Ccl21*) at 4 weeks post-TAC (**Suppl. Table S2**). In contrast, cardiac western blot analyses at 8 weeks revealed increased activation of VEGF-C, as the ratio between mature and immature forms of the growth factor was elevated post-TAC, as compared to sham mice (**Fig. 4a**), while VEGF-D protein levels were not altered (**Suppl. Fig. 3a**). These results are in line with our previous report that cardiac lymphangiogenesis post-TAC in BALB/c mice is essentially mediated by VEGF-C.(*11*) Interestingly, we observed a subpopulation-dependent correlation of cardiac macrophage densities and lymphatic densities post-TAC, with infiltrating macrophages potentially exerting deleterious effects, while tissue-resident macrophages were linked to enhanced lymphangiogenesis (**Suppl. Fig. 3b, c**).

**Figure 4.**
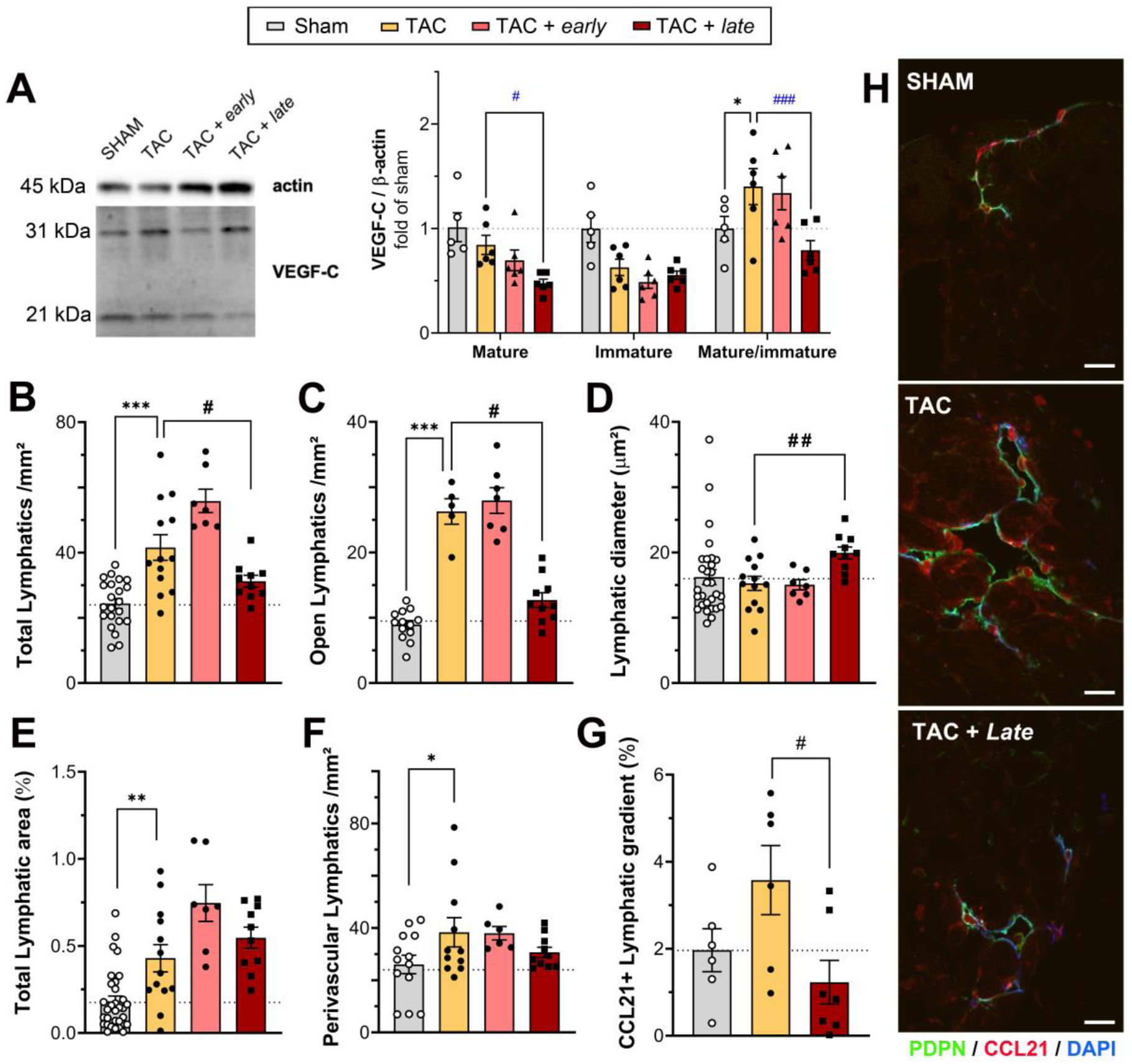
*Late* anti-IL-1β treatment limits TAC-induced lymphangiogenesis. Examples and quantification (**a**) of cardiac VEGF-C protein levels (21 kDa mature vs 31 kDa immature monomeric isoforms) at 8 weeks post-TAC in sham (*n=5;* grey bars w. white circles), control TAC (*n=5;* yellow bars w. black circles), and *early* (*n=5;* red bars w. black circles) vs. *late* (n*=5;* red bars w. black squares) anti-IL-1β-treated TAC mice. Cardiac capillary lymphatic density (**b**), open vessel density (**c**), lymphatic vessel lumen sizes (**d**), lymphatic vascular area (**e**), and perivascular lymphatic density (**f**) evaluated in histological sections at 8 weeks (*n=6- 13* mice per group). Quantification of CCL21 gradients in cardiac lymphatics at 8 weeks (**g**). Examples of cardiac lymphatic chemokine expression (only extracellular signal evaluated to determine extent of gradient) (**h**): Podoplanin (PDPN), *green*; CCL21, *red*, Dapi *blue*. Scalebar 50 µm. Mean ± s.e.m are shown. Groups were compared by two-way ANOVA followed by Dunnett’s multiple comparisons test (**a**), one-way ANOVA followed by Dunnett’s posthoc multiple comparisons test (**b**), non-parametric Kruskal Wallis followed by Dunn’s posthoc (**c-e, g**), or one-way ANOVA with Sidak posthoc (**f**). * p<0.05, **p<0.01, ***p<0.001 *versus* sham, #p<0.05, ## p<0.01, ### p<0.001 *versus* TAC control.

Next, to investigate whether IL-1β could increase VEGF-C production in immune cells, we stimulated bone marrow-derived macrophages for 4h or 24h and analyzed expression levels of *Vegfc* and associated genes encoding proteins involved in VEGF- C maturation, e.g. Adamts3 (*A disintegrin and metalloproteinase with thrombospondin motifs-3*) that cleaves VEGF-C N-terminal propeptide under guidance of Ccbe1 (*Collagen and calcium-binding EGF domain-containing protein-1*)(*30*). We found that IL-1β induced transient upregulation of *Ccbe1*, while *Vegfc* and *Adamts3* levels were increased at both 4h and 24h of stimulation (**Table 2**). Further, whereas IL-1β did not impact the expression of Cathepsin D (*Ctsd*) or proprotein convertases (*Psck5* or *Psck7*), known to modulate VEGF-C maturation(*30*), it did stimulate the expression of *Furin,* an intracellular proprotein convertase that matures pro-VEGF-C by cleaving its C-terminal propeptide. In agreement, western blot analyses of macrophage-conditioned media revealed an increase in mature VEGF-C levels following IL-1β stimulation (**Suppl. Fig. 4**).

**Table 2.** IL-1β-induced gene expression in macrophages in vitro.

| Gene | 4h |  |  | 24h |  |  |
| --- | --- | --- | --- | --- | --- | --- |
| | Control | IL-1β | *p* | Control | IL-1β | *p* |
|  | N=10-14 | N=10-14 |  | N=7 | N=13-14 |  |
| *Adamts3* | 1.0 ± 0.06 | 1.7 ± 0.17 | *** | 1.0 ± 0.06 | 1.9 ± 0.20 | ** |
| *Adamts14* | 1.0 ± 0.07 | 1.2 ± 0.05 | *n.s* | 1.0 ± 0.10 | 1.1 ± 0.10 | *n.s* |
| *Ctsd* | 1.0 ± 0.07 | 1.2 ± 0.14 | *n.s* | 1.0 ± 0.06 | 1.1 ± 0.07 | *n.s* |
| *Ccbe1* | 1.0 ± 0.05 | 1.7 ± 0.08 | *** | 1.0 ± 0.04 | 1.4 ± 0.12 | *n.s* |
| *Furin* | 1.0 ± 0.05 | 1.1 ± 0.06 | *n.s* | 1.0 ± 0.04 | 1.3 ± 0.05 | ** |
| *Pcsk5* | 1.0 ± 0.08 | 0.6 ± 0.08 | ** | 1.0 ± 0.04 | 0.8 ± 0.15 | *n.s* |
| *Pcsk7* | 1.0 ± 0.04 | 1.0 ± 0.04 | *n.s* | 1.0 ± 0.07 | 1.1 ± 0.08 | *n.s* |
| *Vegfc* | 1.0 ± 0.05 | 1.7 ± 0.07 | *** | 1.0 ± 0.05 | 1.3 ± 0.07 | * |

*Adamts*, A Disintegrin And Metalloproteinase with Thrombospondin Motifs; *Ctsd*, Cathepsin D; *Ccbe1*, collagen- and calcium-binding EGF-like domains 1; *Pcsk*, proprotein convertase. Data reported as fold of control (mean ± sem). Samples were compared using Two-way ANOVA (mixed-effects model) followed by Sidak's multiple comparison test. \* $p < 0.05$ ; \*\* $p < 0.001$ ; \*\*\* $p < 0.001$ versus untreated controls.

To investigate potential direct effects of IL-1β on lymphatics, we stimulated cultured murine lymphatic endothelial cells (LECs) with recombinant cytokine for 24h. The cells expressed low levels of the IL1 receptor *Il1r1*, which were further reduced by IL-1β stimulation (**Suppl. Tables S3, S5**). We found that IL-1β increased *Flt4* levels but did not alter *Vegfc, Vegfd, Ctsd* or *Furin* expression in LECs. The expression levels of other proteases involved in VEGF-C maturation (*Pcsk7, Adamts3, Adamts14, Pcsk5, Ccbe1*) were either very low or undetectable in LECs, although IL-1β slightly increased *Ccbe1* expression. Taken together, our findings support a majoritarian cell non-autonomous role of IL-1β in stimulating lymphangiogenesis. Further, our data align with previous reports demonstrating a key role of macrophages in regulation of cardiac lymphangiogenesis post-TAC(*29*) as well as post-MI(*31*).

*Early* anti-IL-1β treatment did not modify cardiac VEGF-C levels or lymphatic densities at 8-weeks post-TAC (**Fig. 4a-e**), nor did it alter cardiac expression of key lymphatic markers and regulators (*Lyve1, Pdpn, Ccl21, Vegfc*, and *Vegfd*) compared to control TAC mice (**Suppl. Table S4**). In contrast, in support of a role of IL-1β in linking LV dilation to lymphangiogenesis, *late* anti-IL-1β, initiated at 4 weeks post-TAC, and prior to LV dilation, potently reduced cardiac mature VEGF-C levels (**Fig. 4a**). This led to reduced lymphatic densities at 8 weeks (**Fig. 4b, c, Suppl. Fig. 5a**) despite pronounced LV dilation in *late*-treated TAC mice (**Fig. 1b, c**). Interestingly, while lymphatic capillary and precollector (open vessel) densities were reduced, lymphatic vessel sizes were increased (**Fig. 4d**). This enlargement of lymphatics (“profile switch” from 5-20 µm size range to 20-50 µm vessel diameters (**Suppl. Fig. 5b**)) was sufficient to partially compensate for the reduction in lymphangiogenesis following anti-IL-1β treatment, and total cardiac lymphatic areas remained unchanged, as compared to TAC controls (**Fig. 4e**). In contrast, coronary perivascular lymphatic density, which increased post-TAC, was not altered by either *early* or *late* anti-IL-1β treatment (**Fig. 4f**).

Although cardiac expression analyses indicated that *late* anti-IL-1β treatment also reduced *Vegfd* levels (**Suppl. Table S4**), western blot analyses did not confirm this (**Suppl. Fig. 3a**). Further, despite the reduced lymphatic density, we found that *late* anti-IL-1β treatment increased cardiac *Lyve1* and *Ccl21* gene expression, as compared to control TAC mice (**Suppl. Table S4**). However, immunohistochemical analyses revealed that Ccl21 lymphatic gradients were not increased, but paradoxically decreased in *late*-treated mice (**Fig. 4g, h**). Notably, reduced Ccl21 gradients may limit lymphatic drainage of Ccr7^+^ immune cells, including macrophages and T cells. However, as noted above, we did not observe any increase in cardiac immune cells at 8 weeks in *late* anti-IL-1β-treated mice compared to TAC controls. Thus, the mechanisms, and functional relevance, of reduced cardiac lymphatic Ccl21 gradients in *late* anti-IL-1β-treated mice remain to be investigated. Moreover, given these discrepancies between cardiac gene and protein expression levels in *late* anti-IL-1β-treated mice, it is possible that inhibition of IL-1β may also have influenced cardiac proteases related to Ccl21 maturation(*32*).

### 4. Early, but not late, anti-IL-1β treatment modulates cardiac fibrosis post-TAC

While IL-1β acting on fibroblasts stimulates fibrosis(*6, 7*), by stimulating lymphangiogenesis it may enhance cardiac lymphatic drainage and thus reduce edema, inflammation, and fibrosis.(*23*) In our experimental study, we unexpectedly found that collagen production, evidenced as cardiac *Col1a1* gene expression, was increased at 4 weeks by *early* anti-IL-1β treatment (**Suppl. Table S2**). Moreover, *early,* but not late, anti-IL-1β treatment increased interstitial fibrosis at 8 weeks, as compared to control TAC mice (**Fig. 5a**). In contrast, neither *early* nor *late* anti-IL-1β treatments altered TAC-induced development of replacement fibrosis (micro-scars) (**Fig. 5b**) or perivascular fibrosis (**Fig. 5c**), as evaluated in small coronary arterioles (16-60 µm diameter) (**Fig. 5d**). In agreement, we recently reported that BALB/c mice were protected against development of TAC-induced perivascular fibrosis, compared to C57Bl/6J mice.(*11*) We proposed that this may be due to expansion of perivascular lymphatics noted only in the former. Thus, perivascular lymphangiogenesis during pressure-overload may prevent fibrosis locally in the perivascular niche. We previously reported a tendency (p=0.1) for increased lymphatic density around coronaries in clinical biopsies from HF patients, especially in the setting of DCM.(*11*) In the current study, we reanalyzed the data focusing on blood vessels of similar size and mural thickness as analyzed in the mouse samples (16-60 µm arterioles), finding again a tendency for an increase in DCM patients (**Fig. 5e**). Histological analyses revealed that this was associated with lower levels of perivascular fibrosis in DCM compared to ischemic HF patients (**Fig. 5f-h**). Previous studies have similarly reported low levels of perivascular fibrosis in DCM patients.(*26*) Investigating the correlation between the extent of coronary perivascular fibrosis and lymphatic densities in humans and mice, we uncovered a negative correlation (**Fig. 5i**), supporting a local role of lymphatics in limiting fibrosis in the perivascular niche. Our findings that anti-IL-1β treatment in mice did not modify perivascular lymphatic densities or perivascular fibrosis post-TAC indicate that this process is independent of IL-1β.

**Figure 5.**
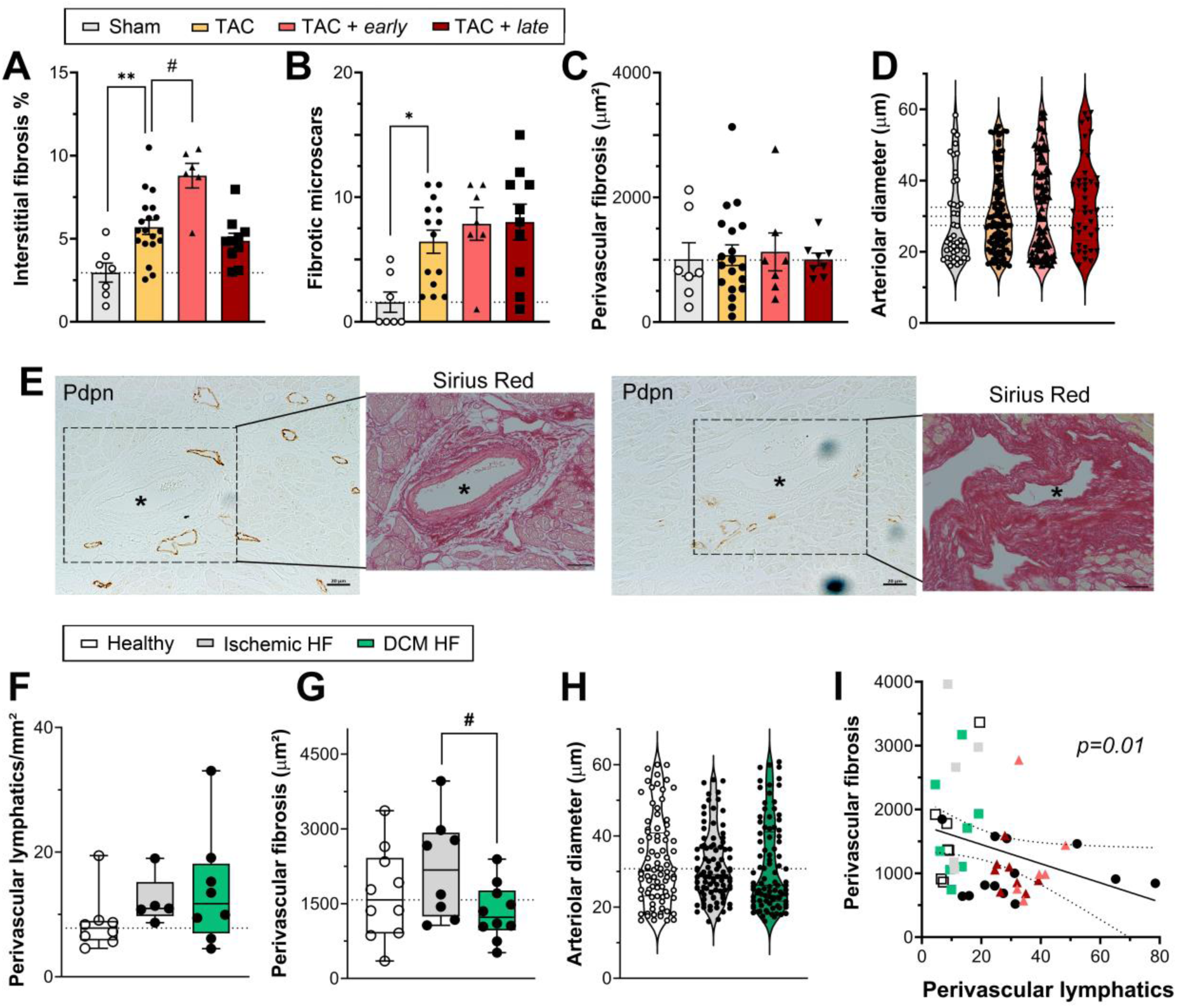
Cardiac perivascular fibrosis post-TAC in mice and HF patients linked to perivascular lymphatics, independently of IL-1β. Evaluation of interstitial fibrosis (**a**) at 8 weeks post-TAC in sham (*n=7;* grey bars w. white circles), control TAC (*n=19;* yellow bars w. black circles), and *early* (*n=6;* red bars w. black circles) vs. *late* (n*=10;* red bars w. black squares) anti-IL-1β-treated TAC mice. Quantification of microscars (**b**) and perivascular fibrosis (**c**) at 8 weeks post-TAC. Arteriolar sizes for evaluation of perivascular fibrosis in mice (**d**). Examples (**e**) of cardiac perivascular lymphatics, stained for PDPN (brown), and perivascular fibrosis in Sirius Red-stained clinical HF samples; scalebar 20 µm. Quantification of perivascular lymphatic densities (**f**, adapted from Heron *et al.*(*11*)) and of perivascular fibrosis (µm²) in healthy humans (*n=8-10, white bar*), ischemic HF patients (*n=5-8, grey bar*), and DCM patients (*n=8-10, green bar*) (**g**). Arteriolar sizes for evaluation of perivascular fibrosis and lymphatic densities in humans (**h**). Correlation of perivascular fibrosis (µm²) and lymphatic density (vessels/mm²) around coronary arteries in humans (*white, grey or green* squares) vs. in TAC-operated mice, either untreated (circles) or anti-IL1b-treated (triangles) (**i**). Non-parametric Spearman rank order test was used to determine correlation (**i**). Mean ± s.e.m are shown (a, b, c), or violin or box plot with median indicated (d, f, g, h). Groups compared by one-way ANOVA followed by Sidak’s multiple comparison test (f), or non-parametric Kruskal Wallis followed by Dunn’s posthoc test (a, b). * p<0.05; ** p<0.01 *versus* sham; # p<0.05 *versus* TAC control or DCM patients.

## Discussion

In this study, we demonstrated that anti-IL-1β treatment with *Gevokuzimab* during pressure-overload did not significantly alter cardiac inflammation at 8 weeks, nor was it sufficient to prevent deleterious cardiac remodeling, fibrosis, or HF development at 8 weeks post-TAC. In contrast, in line with our hypothesis, we found that IL-1β plays a key role in mediating LV dilation-induced cardiac lymphangiogenesis following pressure-overload, occurring between weeks 4-8 post-TAC in BALB/c mice. Indeed, TAC-induced cardiac lymphangiogenesis was reduced only by *late* anti-IL-1β treatment, initiated week 4, but not by *early* anti-IL-1β treatment, discontinued in week 5. Our data further revealed that *late* anti-IL1β did not influence cardiac macrophage levels or *Vegfc* gene expression in the heart. Instead, we found reduced cardiac mature VEGF-C protein levels in *late*-treated mice. Our *in vitro* data support a direct role of IL- 1β in promoting macrophage *Vegf-c* expression and VEGF-C maturation, linked to upregulation of *Furin, Ccbe1*, and *Adamts3*. In addition, IL-1β may also enhance lymphangiogenesis by stimulating LEC expression of *Flt4*. Indeed, although LECs express low levels of *Il1r1*, IL-1β has been shown to enhance LEC NFκB signaling, promoting *Flt4* expression thus sensitizing LECs to autocrine or paracrine VEGF- C(*33*).

The finding of indirect pro-lymphangiogenic effects of IL-1β are reminiscent of findings reported in mice with tracheal *Il1b*-overexpression.(*21*) In this setting, blockage of VEGF-C/-D signaling, with soluble VEGFR-3, sufficed to inhibit IL-1β-induced tracheal lymphangiogenesis, demonstrating its dependency on the VEGFR-3 axis. The authors further showed that IL-1β stimulated tissue-recruitment of macrophages, and increased their release of VEGF-C. In contrast, we found a negative correlation between infiltrating macrophages and lymphatic densities in the heart post-TAC. Thus, in our study, cardiac IL-1β likely did not stimulate lymphangiogenesis post-TAC by promoting the recruitment of macrophages. Indeed, in line with a previous report in C57BL6 mice(*29*), we found a weak, but positive, correlation between densities of tissue-resident cardiac macrophages and lymphatics. However, IL-1β does not appear as a major regulator of LYVE1^+^ tissue-resident cardiac macrophage expansion, as neither *early* nor *late* phase blockage of IL-1β post-TAC altered densities of this subpopulation. Our *in vitro* data indicated that IL-1β may influence macrophage activity, rather than numbers, promoting VEGF-C maturation. Indeed, we demonstrate that IL-1β stimulation of macrophages, but not LECs, led to upregulation of *Vegfc* and its proteases, including *Furin* and *Adamts3*, essential for VEGF-C maturation. Of note, recent cardiac single-cell RNA sequencing (scRNAseq) analyses further suggested that IL-1β may regulate extracellular matrix remodeling by increasing cardiac fibroblast expression of endogenous protease inhibitors, including TIMPs(*34, 35*).

In addition to stimulating VEGF-C-dependent macrophage-induced lymph- angiogenesis post-TAC, our study suggests that IL-1β may influence lymphatic immune cell recruitment by enhancing Ccl21 release in LECs. Indeed, we found reduced cardiac lymphatic Ccl21 chemokine gradients in *late* anti-IL-1β-treated mice. This may limit immune cell drainage from the heart, thus slowing inflammatory resolution, as Ccl21 is the main lymphatic chemokine, and its receptor, Ccr7, is expressed in both lymphoid and myeloid cells (*36*)^-^(*37*). Our *in silico* analysis of recently published scRNAseq data, from cardiac-infiltrating immune cells in C57BL6/J mice post-TAC(*38*) or post-MI(*25*), indicate that both B and T cells have elevated *Ccr7* expression in the heart. Thus, reduced lymphatic Ccl21 levels, together with reduced lymphatic density following inhibition of IL-1β, could potentially increase cardiac T cell retention. Our findings, however, did not confirm increased cardiac T cell levels in anti-IL-1β-treated TAC mice.

We also expected that reducing IL-1β signaling, and thus limiting cardiac lymphangiogenesis, during pressure-overload may alter cardiac levels of infiltrating macrophages. While we found no differences in cardiac-resident CD68^+^/Lyve-1^+^ macrophages, known to be cardioprotective post-TAC(*29*), in either *early* or *late*- treated groups, there was a transient increase in CD68^+^/Lyve-1^neg^/CD206^+^ macrophages, a subpopulation classified as alternatively activated(*39*), in *early* anti-IL-1β treated mice. Importantly, in *late*-treated mice, where IL-1β-mediated- lymphangiogenesis was inhibited, the absence of increased cardiac immune cell levels indirectly suggests cardiac lymphatic dysfunction post-TAC. Indeed, we previously reported that the expanded cardiac lymphatic network in BALB/c mice post-TAC was inefficient in resolving cardiac edema and inflammation, and thus failed to prevent interstitial fibrosis, LV dilation and HF development.(*11*)

We were also surprised by the absence of functional cardiac effects of anti-IL-1β treatment at 8 weeks post-TAC, although *early* anti-IL-1β treatment transiently delayed LV dilation. These results contrast with previous studies demonstrating that *Nlrp3* gene deletion (reducing IL-1β production), or early anti-IL-1β antibody therapy, accelerated cardiac dysfunction post-TAC in C57Bl/6J mice, despite reducing cardiac hypertrophy, macrophage infiltration, and fibrosis.(*9, 10*) These studies suggested that prevention, through blocking of IL1β, of adaptive hypertrophy (induced by low levels of IL-1β in C57Bl/6J mice) promoted LV dilation and dysfunction post-TAC. Additionally, the authors suggested insufficient angiogenesis matching cardiac hypertrophy as a driver of accelerated cardiac dysfunction in *Nlrp3*-deficient or anti-IL-1β-treated mice(*9, 10*). In contrast, in BALB/c mice, pressure-overload does not lead to extensive cardiac hypertrophy, thus lessening the role of cardiac angiogenesis and the functional impact of potential anti-hypertrophic effects of anti-IL-1β treatment. Indeed, our previous studies revealed maintenance, but not expansion, of blood vascular networks post-TAC in BALB/c mice.(*11*) *Of note, Gevokizumab* decreases the binding affinity of IL- 1β to IL-1R, resulting in a 30-fold reduction of IL-1β signaling, but not a complete block of IL-1β signaling, different from *Canakinumab* and *Anakinra*.(*40*) We expect that in our study *early* anti-IL-1β treatment may have reduced, but not completely prevented, cardiac IL-1β signaling, thus maintaining the moderate but beneficial compensatory hypertrophy(*5*), while transiently blocking the detrimental direct (negative inotropic(*4*)) effects of IL-1β on cardiomyocytes. It is possible that *early* and *prolonged* anti-IL-1β treatment would have been required in our study to reach durable benefit on cardiac function. However, long-term treatment, including during LV dilation, would have reduced cardiac lymphangiogenesis, thus potentially confounding beneficial direct cardiac effects of IL-1β blockage. Of note, the risk for an immune response against the therapeutic antibody prevents long-term treatment with *Gevokizumab* in mice.

Other pro-inflammatory cytokines induced by cardiac wall stretch, such as IL-6(*4*), may also potentially counteract reduced IL-1β signaling and thus contribute to the DCM-like phenotype post-TAC in BALB/c mice. IL-6 has been shown to drive both cardiac fibrosis and pathological hypertrophy following pressure-overload in C57BL6 mice.(*41*) We found that anti-IL-1β treatment did not reduce cardiac *Il6* expression, which may have contributed to the lack of beneficial effects on cardiac fibrosis and HF development in both *early-* and *late*-treated mice. Importantly, the lack of cardiac benefit of *late* anti-IL-1β treatment in our study suggests that in patients with chronic pressure-overload IL-1β-blocking therapy may be inefficient, and potentially deleterious, by reducing cardiac lymphatic density if initiated during LV dilation.

Intriguingly, the beneficial perivascular lymphangiogenesis occurring post-TAC, which seems to protect against perivascular fibrosis, was not influenced by anti-IL-1β treatment. In agreement, recent scRNAseq analyses coupled to spatial transcriptomics revealed that perivascular fibroblasts, different from Postn^+^ matrifibroblasts, have little interaction with cardiac-infiltrating IL-1β-producing Ccr2^+^ myeloid cells(*34*). Further studies are needed to identify the triggers of lymphangiogenesis in the perivascular niche in the heart and to unravel how these lymphatics limit perivascular fibrosis during HF development. Notably, a histological marker of edema would be required to investigate if these expanded lymphatics surrounding coronary arterioles suffice to locally restore perivascular pressure-gradients caused by excessive blood vascular leakage occurring during pressure-overload(*11*).

Taken together, our study provides, for the first time, a molecular mechanism linking LV dilation to lymphatic remodeling following pressure-overload (**Fig. 6**). Moreover, our data indicate that IL-1β may influence not only lymphatic expansion, but also macrophage and lymphatic cross-talk. Intriguingly, the absence of deleterious cardiac effects following inhibition of lymphangiogenesis by *late* anti-IL-1β treatment is in support of our previous finding that the expanded lymphatic network in BALB/c mice post-TAC is dysfunctional and/or immature(*11*). In contrast, we and other have demonstrated that inhibition of lymphangiogenesis during pressure-overload in C57BL6/J mice and Wistar rats, by *Flt4* deletion or anti-VEGFR-3 treatment, aggravates cardiac inflammation and dysfunction.(*11, 17, 42, 43*) Further investigations are needed to determine why IL-1β-mediated lymphangiogenesis during pressure-overload leads to aberrant hyper-sprouting and dysfunctional cardiac lymphatics in BALB/c mice. Molecular analyses are underway to determine how expanded cardiac lymphatics post-TAC differ from healthy lymphatics in BALB/c mice, to identify new targets to treat lymphatic dysfunction in HF(*44*).

**Figure 6.**
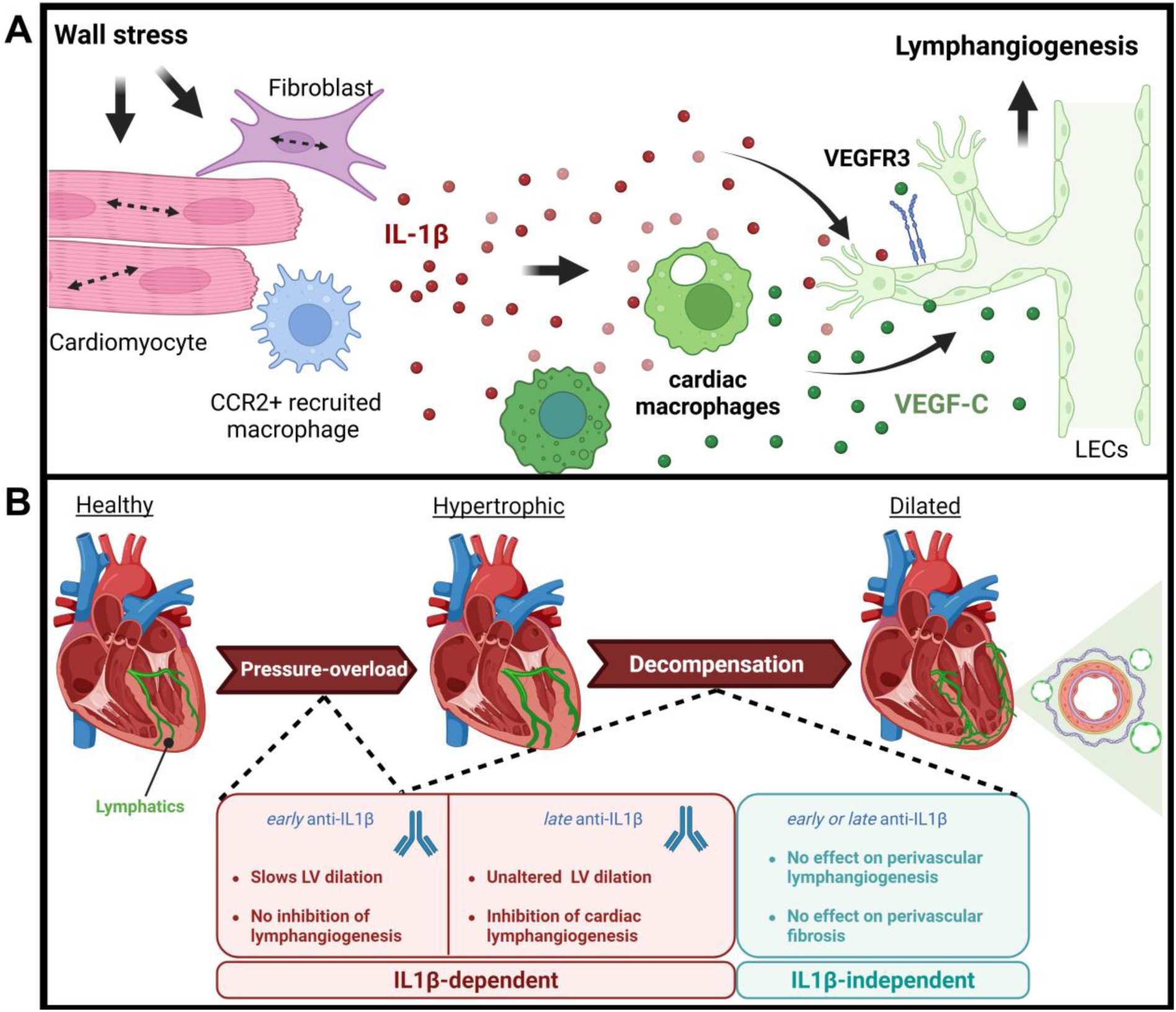
Schematic view of cardiac impact of IL-1β during pressure-overload. In response to increased LV wall stress (**a**), cardiac cells, including cardiomyocytes, fibroblasts and recruited CCR2^+^ macrophages, release IL-1β. This stimulates both production and maturation of VEGF-C by cardiac macrophages. In parallel, IL-1β acts on LECs to increase VEGFR3 (*Flt4*) expression. Together, these changes lead to stimulation of cardiac lymphangiogenesis. In cardiac pathology (**b**), LV dilation triggers IL-1β-dependent macrophage-mediated stimulation of cardiac lymphangiogenesis. However, in the setting of cardiac inflammation, the expanded lymphatic network remains immature and/or dysfunctional, and fails to limit accumulation of immune cells, edema, and interstitial fibrosis. In contrast, perivascular lymphangiogenesis, triggered in an IL-1β-independent manner during LV dilation, may limit perivascular fibrosis. Antibody-mediated blockage of IL-1β in *early* phase post-TAC in BALB/c mice, before the onset of lymphangiogenesis, transiently limits LV dilation. In contrast, *late* anti*-*IL-1β treatment, initiated during LV dilation, blocks cardiac lymphangiogenesis, but does not alter cardiac remodelling or dysfunction. Created with BioRender.com.

As for the clinical outlook, a recent case-report in a stage III HF patient, secondary to idiopathic inflammatory DCM, demonstrated rapid improvement of LV function following *Anakinra* treatment, associated with reduced IL-6 and BNP levels.(*45*) We speculate that this cardiac benefit of IL-1β inhibition in an already severely dilated heart may be related to the reduction of negative inotropic effects of the cytokine. In our study, we cannot exclude that the observed inhibition of lymphangiogenesis may have contributed to the lack of functional cardiac benefit of anti-IL-1β in *late-*treated mice.

Thus, anti-IL-1β may have a dual therapeutic window where targeting of IL-1β in the early stage, before LV dilation, or alternatively at later stages of the disease, when LV dilation and cardiac lymphangiogenesis are well-established, may be required to obtain clinical benefit in HF patients.

## Author contributions

C.H, O.L, T.L and A.D performed and analyzed immunohistochemistry and histology. Echocardiography was performed and analyzed by T.L and O.L; microgravimetry was carried out by C.H and T.L; M.B and A.Z designed and carried out *in vitro* macrophage assays and expression analyses; M.V carried out surgical mouse model; C.H, T.L and O.L carried-out cardiac gene expression analyses; O.L and C.V carried out *in vitro* LEC assays and expression analyses; C.H and C.V performed protein analysis by ELISA and Western Blot; JB.M managed the biobank at Bichat Hospital and contributed clinical data for this study; P.M, A.Z, and V.T participated in study design; A.Z contributed critical comments on the manuscript draft; C.H. and E.B designed the study, analyzed results, and prepared the manuscript draft. All authors approved of the final version of the manuscript.

## Acknowledgments

We thank Pr Arantxa González Miqueo, (FIMA, Pamplona, Spain) for helpful discussion on fibrosis; and Mrs Saida Morchid Azhar, Ms Nousayba Ouattara, and Ms Héloïse Quesnel (all Normandy University, Rouen, France) for technical assistance with Western blot, histological analysis, and RTqPCR, respectively.

## Funding

C.H, T.L, and C.V were funded by fellowships from the Normandie Doctoral School (EdNBISE). O.L was co-funded by a fellowship “Allocation 50% Normandie Recherche. This work was supported by the Fondation pour la Recherche Médicale (FRM) [*FDT202204014960*] to C.H and [FDT202404018270] to T.L. V.T was funded by the Chair of Excellence program “*Lymphcosign*” from the Normandy Region (RIN Recherche). The research leading to these results benefitted from funding (E.B) from the ERA-CVD Research Programme, which is a transnational R&D programme jointly funded by national funding organisations [*ANR-16-ECVD-0004*] within the framework of the ERA-NET ERA-CVD. This work is part of the FHU-CARNAVAL supported by a grant from the GCS G4 and labeled by AVIESAN. This work was further supported by a joint grant (E.B, A.Z) from the Agence Nationale de la Recherche (ANR) with the Deutsche ForschungsGemeinschaft (DFG) for the project “CITE-LYMPH” [*ANR-22- CE92-0040-001*; DFG project number 505700170], and DFG projects 453989101 – SFB1525 and 324392634 - TRR221 (A.Z.). The project also benefitted from generalized institutional funds (Inserm U1096 laboratory) from French Inserm, Rouen University (BQRI 2023) and the Normandy Region (CPER 2021, 2022) together with the EU-Normandy region co-funding (2019 7D MICROSCOPY) “*L’Europe s’engage en Normandie avec le Fonds Européen de Développement Régional*”. We acknowledge France-BioImaging infrastructure (https://ror.org/01y7vt929) supported by the French National Research Agency (ANR-24-INBS-0005 FBI BIOGEN).

## Conflict of Interest

The authors declare that they have no conflict of interest.

## Data Availability

The original data included in this study has been deposited in GEO (GSE290146, *Effect of interleukin-1beta on confluent murine lymphatic endothelial cells*).

## Notes

### Competing Interest Statement

The authors have declared no competing interest.

### Summary of Updates

Cardiac macrophage evaluation performed at 4 weeks post-TAC RNAseq performed in murine LECs stimulated with IL1beta More detailed description of methods Updated figures & references

