## Supplementary material for "Blocking Interleukin-1β transiently limits left ventricular dilation and reduces cardiac lymphangiogenesis during pressure-overload in mice": Suppl Figures, Tables S1-S4, and Methods

### Supplementary methods & materials

**Suppl. Data**

Suppl. Figure S1 Echocardiographic and morphometric analyses at 8 weeks

Suppl. Figure S2 Cardiac cytokine expression at 8 weeks post-TAC

Suppl. Figure S3 Cardiac VEGF-D levels and lymphatic size profiles post-TAC

Suppl. Figure S4 VEGF-C maturation induced by IL-1β in macrophages

Suppl. Figure S5 Cardiac lymphatic network remodelling post-TAC

Suppl. Table S1 Cardiac function evaluated by echocardiography at 6-weeks post-TAC

Suppl. Table S2 Cardiac gene expression at 4 weeks post-TAC

Suppl. Table S3 LEC gene expression after 24h IL1β stimulation *in vitro*

Suppl. Table S4 Cardiac gene expression at 8 weeks post-TAC

**Suppl. Table S5** List of DEGs induced by IL1β stimulation of mouse LECs

**Suppl. Methods**

Suppl. Figure S6 perivascular image analysis

Suppl. Table S5 antibodies used for immunohistochemistry in mice

Suppl. Table S6 list of primers used in mouse cardiac samples for qPCR

Suppl. Table S7 list of primers used in mouse macrophages for qPCR analysis

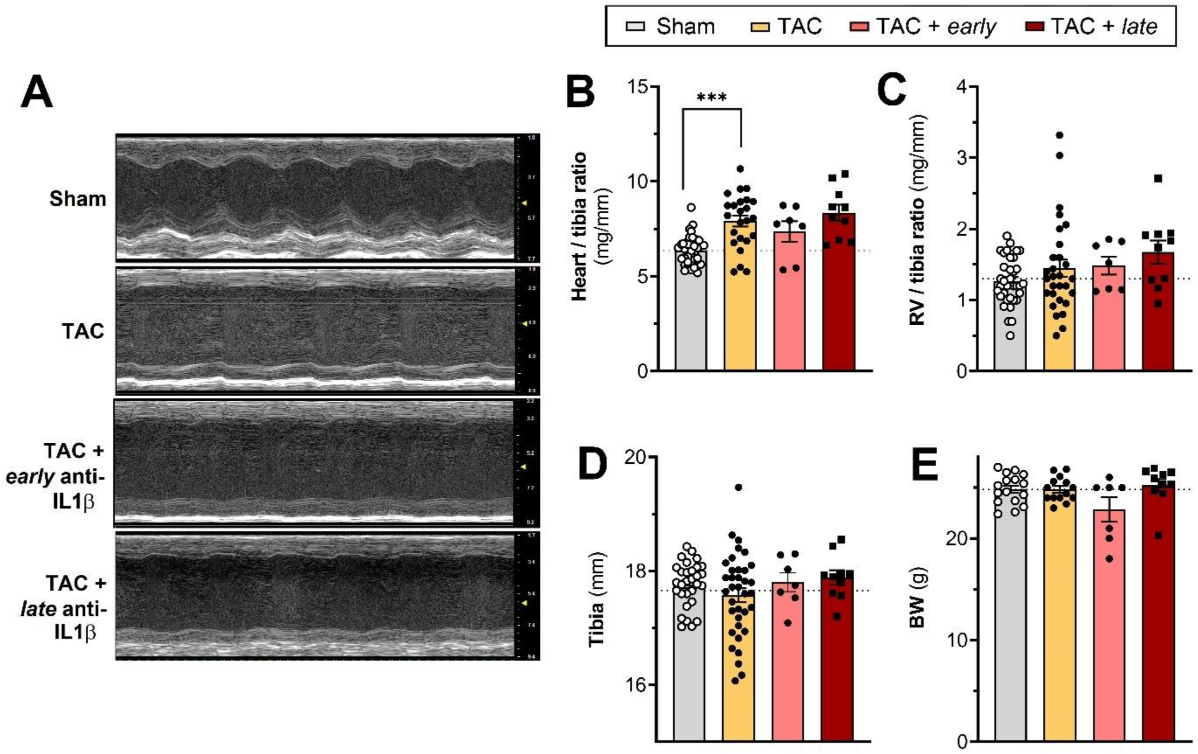


#### Suppl. Fig. 1- Echocardiographic and morphometric analyses at 8 weeks

Examples of echocardiography short-axis views (**a**). Assessment of cardiac (**b**) and right ventricular (RV) weights (**c**), normalized to tibia lengths (**d**), and body weight (**e**) at 8 weeks in sham (*n=16-30;* grey bars w. white circles), control TAC (*n=10-36;* yellow bars w. black circles), and *early* (*n=7;* red bars w. black circles) vs. *late* (n=*10;* red bars w. black squares) anti-IL-1β-treated TAC mice. Data reported as mean ± sem. Groups were compared by one-way ANOVA followed by Sidak's multiple comparison test *** p<0.001 *versus* sham.

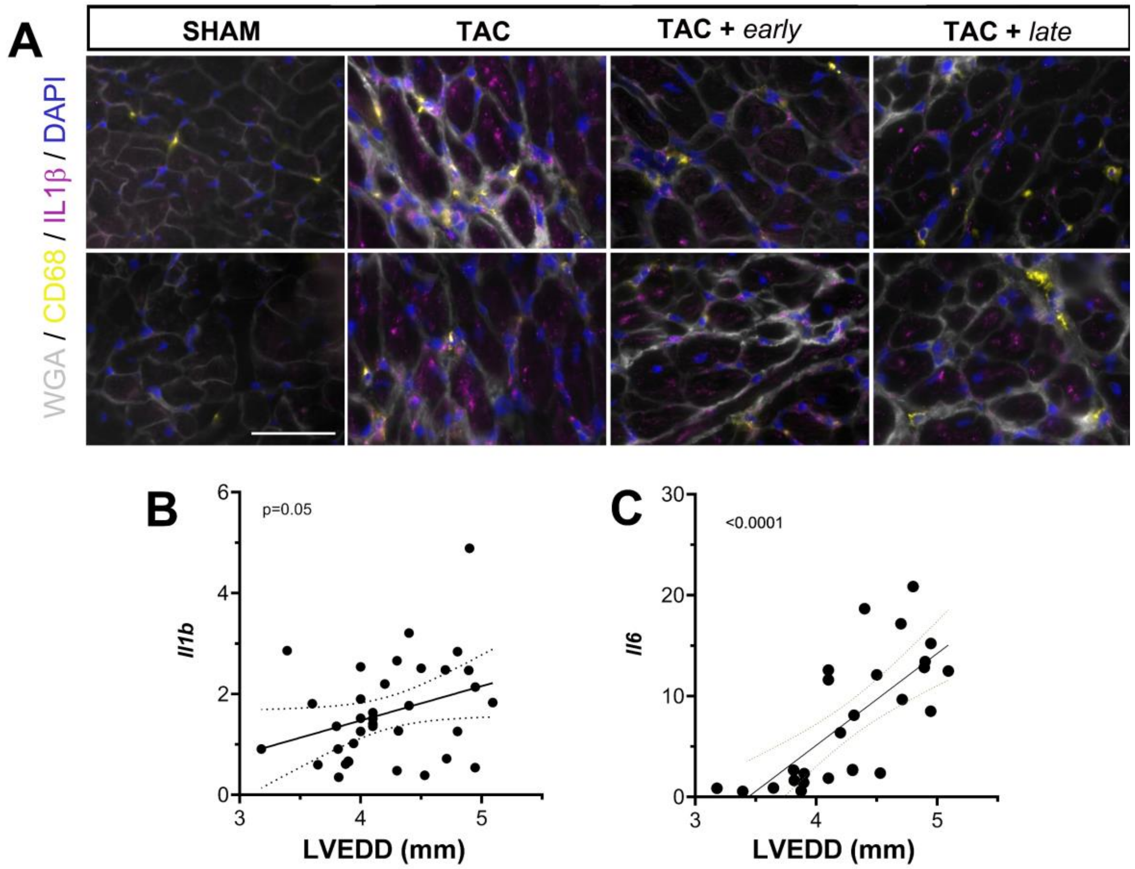


#### Suppl. Fig. 2 -Cardiac cytokine expression at 8 weeks post-TAC

Immunohistochemical analyses of cardiac IL-1β expression in the heart at 8 weeks in sham, control TAC, and *early* vs. *late* anti-IL-1β-treated TAC mice (**a**). Wga, *grey*; macrophages (CD68), *yellow*; IL-1β, *magenta*; and Dapi, *blue*. Scale bar 50 µm. Correlation of cardiac gene expression levels (*fold of sham*) of *Il1b* (**b**), and *Il6* (**c**) with the degree of LV dilation (LV EDD) at 8 weeks post-TAC (*n=6* sham; *n=8* TAC; *n=7* TAC + early anti-IL1β; *n=6* TAC + late anti-IL1β). Non-parametric Spearman rank order tests were used for evaluating correlations.

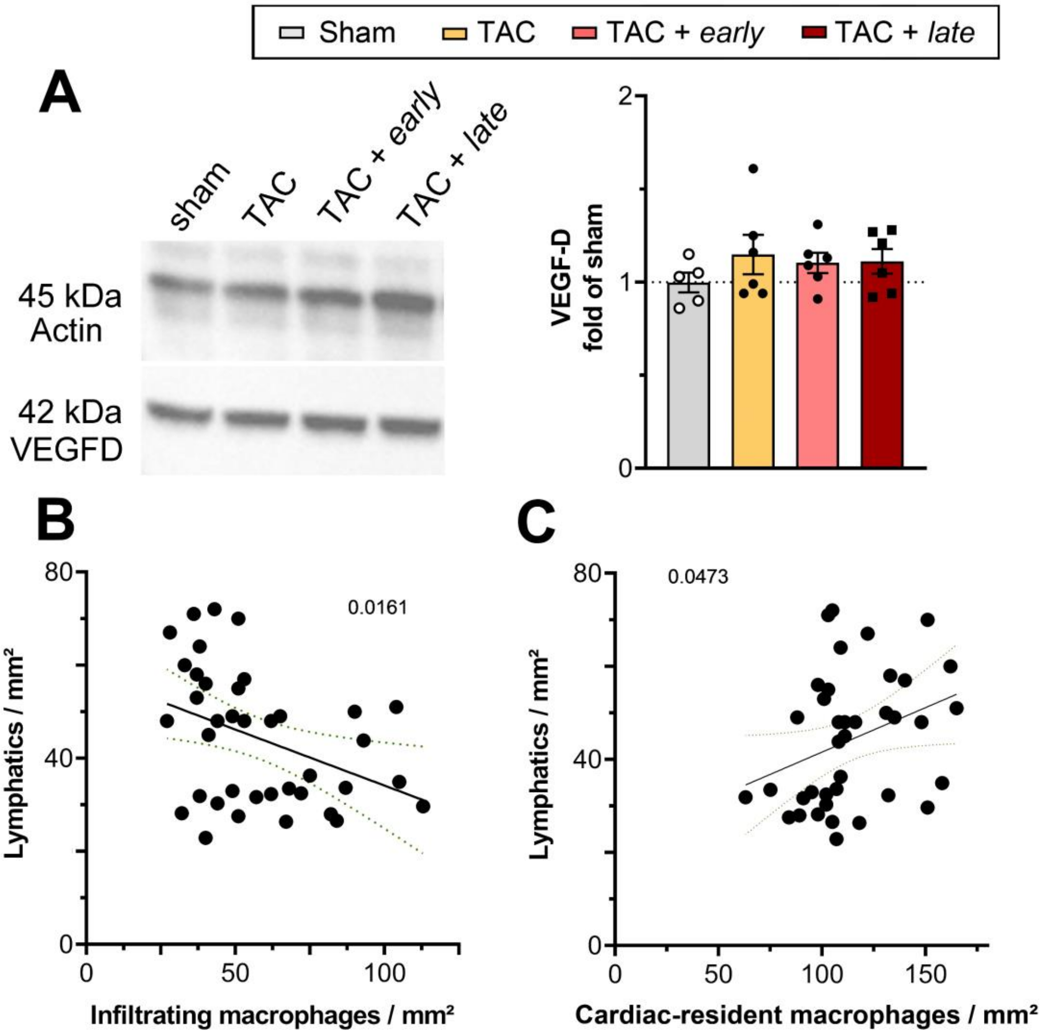


#### Suppl. Fig. 3 -Cardiac VEGF-D levels and lymphatic size profiles post-TAC

Western blot examples and quantification of cardiac VEGF-D (*42 kDa*) protein levels at 8 weeks in sham (*n=5;* grey bars w. white circles), control TAC (*n=5;* yellow bars w. black circles), and *early* (*n=5;* red bars w. black circles) vs. *late* (n=*5;* red bars w. black squares) anti-IL-1β-treated TAC mice as compared to actin (*45 kDa*) (**a**). Correlation between $CD68^{+}$ cardiac-infiltrating ($LYVE1^{neg}$) macrophages (**b**) or resident $LYVE1^{+}$ macrophages (**c**) versus lymphatic densities post-TAC (*n=7-10* mice per group). Data reported as mean ± sem. Non-parametric Spearman rank order tests were used for evaluating correlations.

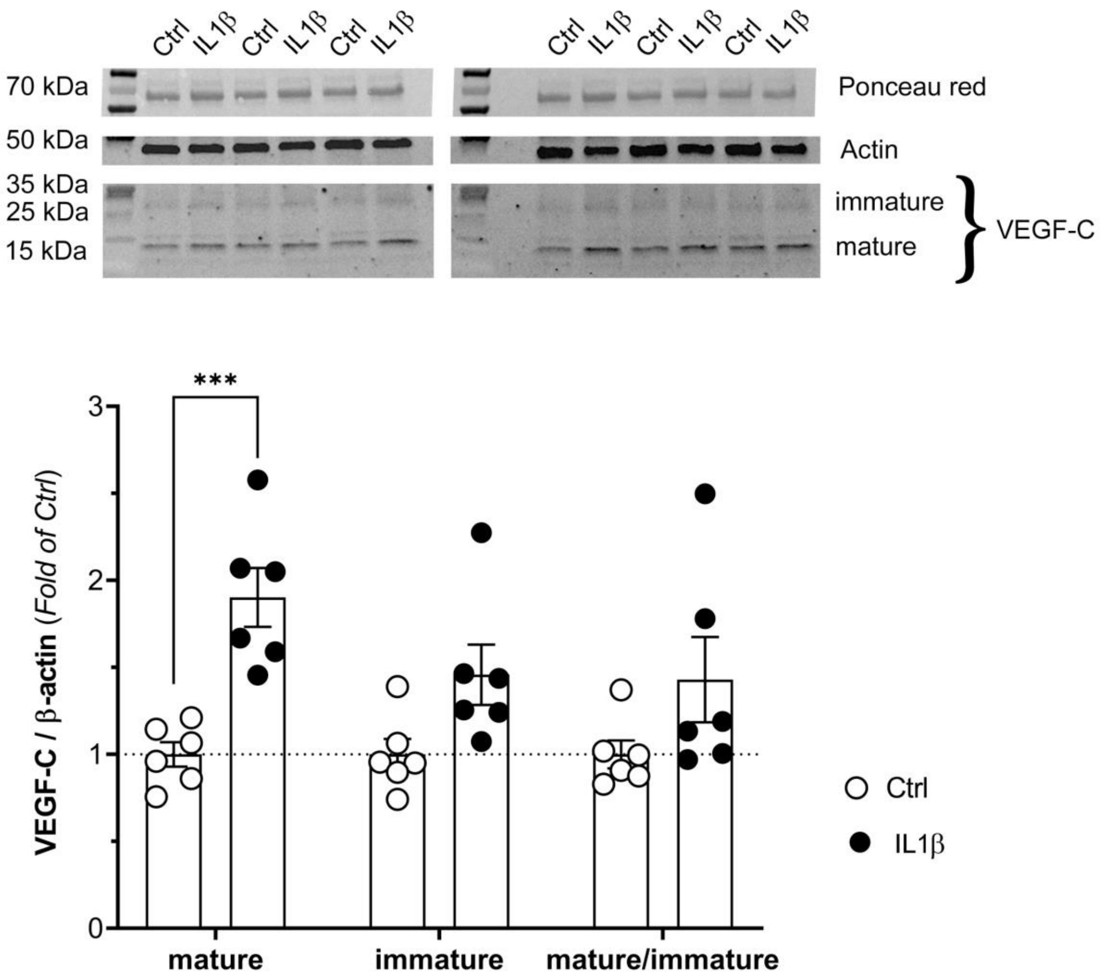


#### Suppl. Fig 4 – VEGF-C maturation induced by IL-1β in macrophages

Examples and quantification of Western blot analysis of VEGF-C levels in macrophage-conditioned media at 24h of culture in control condition (*open circles, n=6*) or after stimulation with IL-1β (*closed circles, n=6*). Immature (*31 kDa*), and mature (*15-21 kDa*) VEGF-C levels were quantified and normalized to actin. Ponceau red loading-control is also shown. Data is reported as mean ± sem (fold of control) of averages of 3 separate experiments. Groups were compared by two-way ANOVA followed by Sidak's multiple comparison test. *** $p<0.001$ *versus* Ctrl.

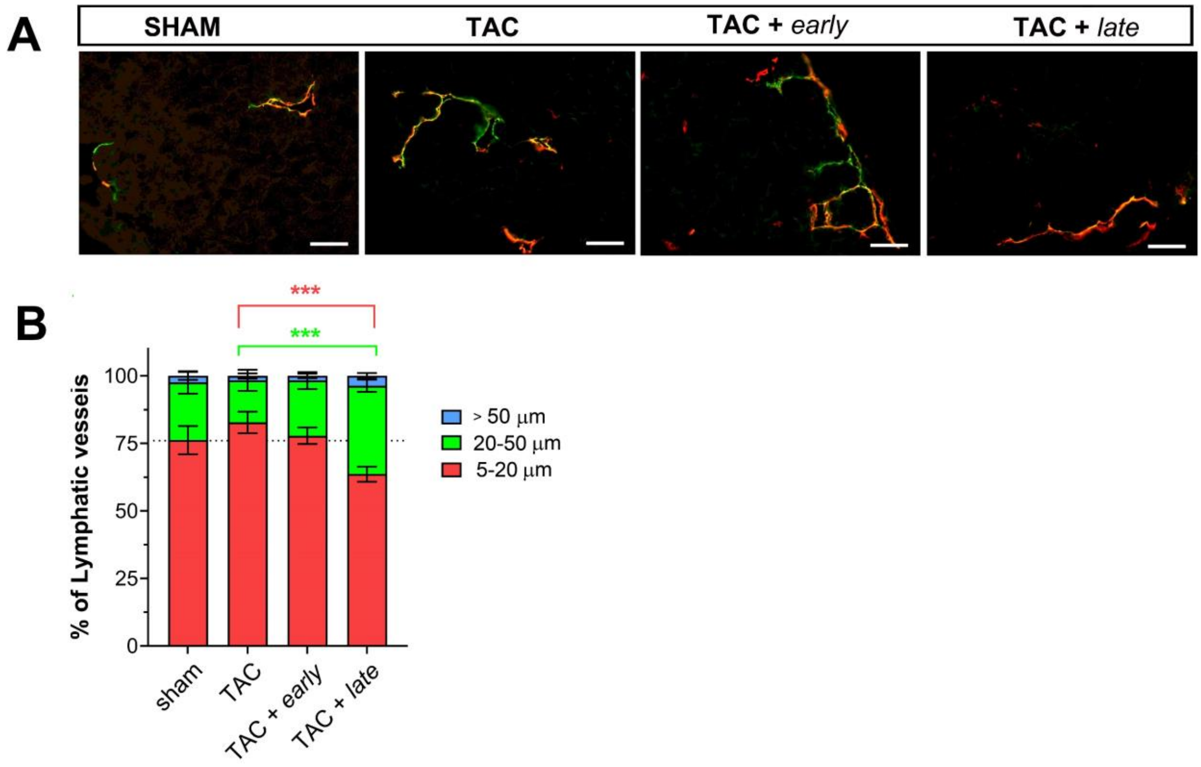


#### Suppl. Fig. 5 -Cardiac lymphatic network remodelling post-TAC

Examples of cardiac lymphatics (**a**) at 8 weeks post-TAC visualized using a combination of LYVE1 (*red*) and Podoplanin (Pdpn, *green*) in cardiac sections. Scalebar 50 µm. Evaluation of lymphatic lumen size profiles at 8 weeks post-TAC (**b**). On average 39 vessels were analyzed per mouse (*n=10* mouse/group). Groups were compared by two-way ANOVA followed by Sidak’s multiple comparison test. *** $p<0.001$ *versus* TAC.

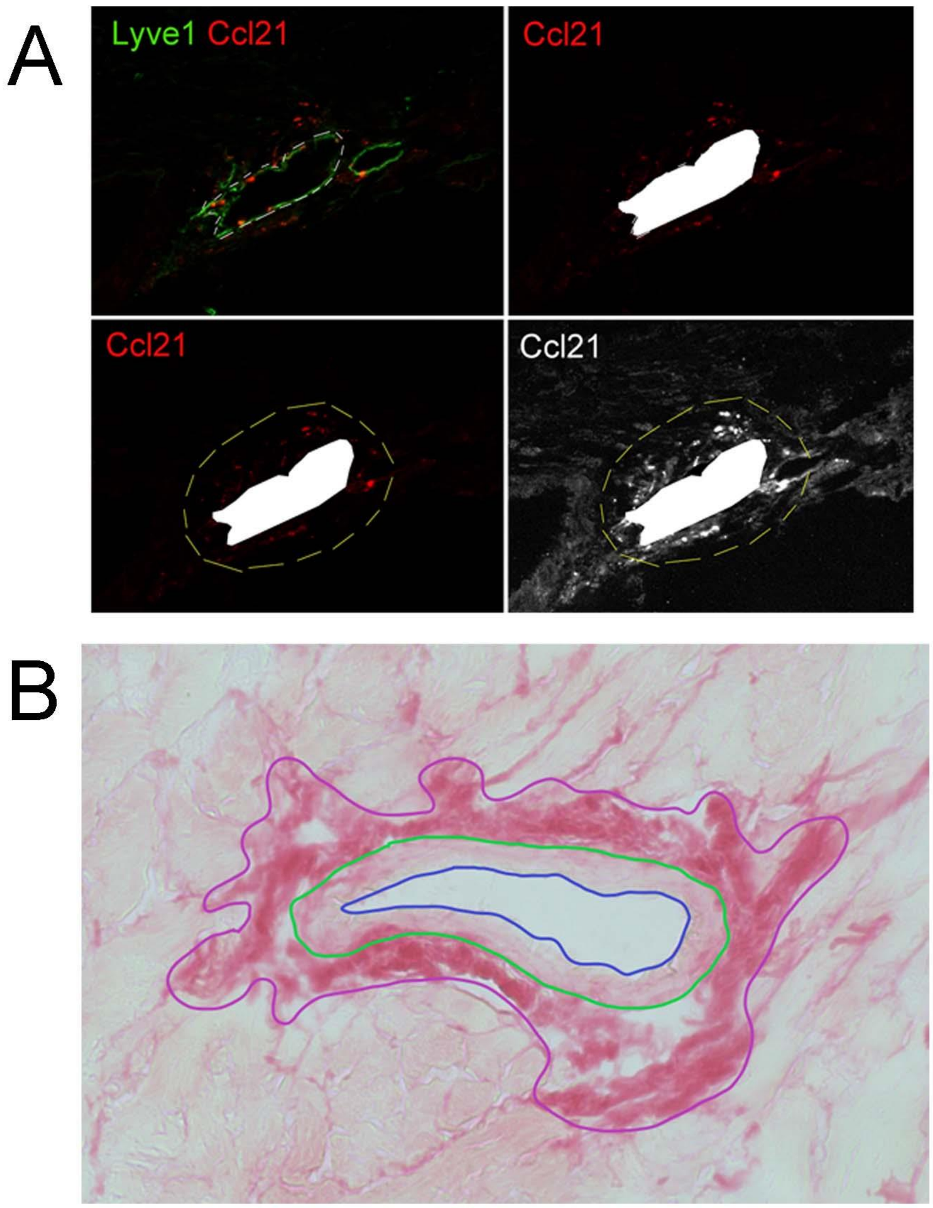


#### Suppl. Fig. 6 -Perivascular image analysis

Examples of cardiac lymphatic Ccl21 gradient evaluation (**a**) performed using ImageJ. White dashed line outlines vessel, yellow dashed line outlines proximal lymphatic area analyzed for Ccl21 positivity. Bottom right quadrant illustrates the threshold set to 15 levels of grey, used to calculate % area surrounding a lymphatic vessel positive for the chemokines. Examples of evaluation of perivascular fibrosis in human and mouse cardiac samples (**b**). ImageJ was used to assess arteriolar sizes (lumen area) [blue contour], muscular arteriolar area [green contour], and the perivascular fibrotic area [purple contour], stained red by Picrosirus dye.

#### Suppl. Table 1 – Cardiac function evaluated by echocardiography at 6-weeks post-TAC

| parameter | | Sham | | TAC | TAC + *early* | | TAC + *late* | |
|---|---|---|---|---|---|---|---|---|
| | | *N=26* | *p* | *N=33* | *N=7* | *p* | *N=10* | *p* |
| **HR** | *bpm* | **355**±9 | *n.s* | **382**±9 | **443**±31 | *n.s* | **391**±14 | *n.s* |
| **AWT ED** | *mm* | **0.8**±0.03 | ** | **0.9**±0.02 | **0.8**±0.07 | *n.s* | **0.9**±0.03 | *n.s* |
| **AWT ES** | *mm* | **1.2**±0.05 | *n.s* | **1.2**±0.04 | **1.21**±0.1 | *n.s* | **1.2**±0.05 | *n.s* |
| **PWT ED** | *mm* | **0.8**±0.04 | *n.s* | **0.9**±0.03 | **0.8**±0.04 | *n.s* | **0.8**±0.04 | *n.s* |
| **PWT ES** | *mm* | **1.1**±0.06 | *n.s* | **1.1**±0.03 | **1.1**±0.04 | *n.s* | **1.0**±0.05 | *n.s* |
| **LVDD** | *mm* | **3.7**±0.06 | *** | **4.3**±0.06 | **3.9**±0.2 | *n.s* | **4.4**±0.1 | *n.s* |
| **LVSD** | *mm* | **2.5**±0.1 | *** | **3.6**±0.08 | **2.9**±0.2 | * | **3.7**±0.1 | *n.s* |
| **FS** | *%* | **33**±2 | *** | **18**±1.1 | **26**±3 | * | **16**±1.8 | *n.s* |
| **VTI** | *cm* | **1.9**±0.1 | *n.s* | **2.0**±0.2 | **2.3**±0.1 | *n.s* | **1.8**±0.1 | *n.s* |
| **SV** | *µL* | **56**±3 | *n.s* | **60**±7 | **59**±5 | *n.s* | **47**±5 | *n.s* |
| **CO** | *mL/min* | **20**±1 | *n.s* | **23**±3 | **26**±3 | *n.s* | **18**±2 | *n.s* |
| **EDV** | *µL* | **60**±3 | *** | **77**±4 | **35**±6 | *** | **89**±4 | *n.s* |
| **ESV** | *µL* | **24**±2 | *** | **60**±4 | **67**±7 | *n.s* | **61**±5 | *n.s* |
| **EF** | *%* | **62**±3 | *** | **39**±2 | **51**±4 | *n.s* | **34**±3 | *n.s* |

*BW*, body weight; *HR*, heart rate; *AWT ED*, anterior wall thickness end-diastole; *AWT ES*, anterior wall thickness end-systole; *PWT ED*, posterior wall thickness end-diastole; *PWT ES*, posterior wall thickness end-systole; *LVDD*, left ventricular diastolic diameter; LVSD, left ventricular systolic diameter; *FS*, fractional shortening: *VTI*, velocity time integral; *SV*, stroke volume; *CO*, cardiac output; *EDV*, end diastolic volume; *ESV* end systolic volume; *EF*, ejection fraction. Data reported as mean ± sem. All samples were compared using Two-way ANOVA followed by Sidak’s posthoc test. * p<0.05; ** p<0,001; *** p<0.001 *versus* TAC controls.

#### Suppl. Table 2 – Cardiac gene expression at 4 weeks post-TAC

| *gene* | Sham | | TAC | TAC + *early* | |
|---|---|---|---|---|---|
| | *N=10* | *p* | *N=9-10* | *N=8-9* | *p* |
| ***Ccl21*** | **1.00**±0.07 | *n.s* | **1.20**±0.13 | **1.34**±0.09 | *n.s* |
| ***Col1a1*** | **1.00**±0.08 | *** | **2.29**±0.25 | **2.94**±0.14 | ** |
| ***Flt4*** | **1.00**±0.07 | *n.s* | **1.02**±0.06 | **1.03**±0.03 | *n.s* |
| ***Lyve1*** | **1.00**±0.06 | *n.s* | **1.10**±0.11 | **1.04**±0.07 | *n.s* |
| ***Pdpn*** | **1.00**±0.07 | *n.s* | **1.02**±0.06 | **1.03**±0.03 | *n.s* |
| ***Tgfb*** | **1.00**±0.07 | *n.s* | **1.35**±0.11 | **1.55**±0.09 | *n.s* |
| ***Vegfc*** | **1.00**±0.05 | *** | **1.46**±0.13 | **1.70**±0.12 | *n.s* |

Data reported as fold of sham (mean ± sem). All samples were compared using Two-way ANOVA followed by Sidak’s posthoc test. * p<0.05; ** p<0,001; *** p<0.001 *versus* TAC controls.

#### Suppl. Table 3 –LEC gene expression after 24h IL1β stimulation *in vitro*

| gene | Control N=3 | Control p | IL-1β N=3 | FC |
|---|---|---|---|---|
| *Adamts3* | N.D | - | N.D | - |
| *Adamts14* | N.D | - | N.D | - |
| *Ctsd /CatD* | 1133±19 | *n.s* | 1103±27 | 0.97 |
| *Ccbe1* | 0.90±0.07 | **p=0.017** | 1.31±0.08 | **1.46** |
| *Flt4* | 34±0.6 | **p=4.7E$^{-26}$** | 50±0.3 | **1.47** |
| *Furin* | 68±1.3 | *n.s* | 59±0.7 | 0.86 |
| *Il1r1* | 14±0.6 | **p=7.1E$^{-05}$** | 11±0.3 | **0.77** |
| *Pcsk5* | 0.01±0.00 | - | N.D | 0.80 |
| *Pcsk7* | 7.1±0.1 | *n.s* | 7.3±0.2 | 1.02 |
| *Vegfc* | 0.46±0.04 | *n.s* | 0.66±0.04 | 1.42 |
| *Vegfd* | 0.54±0.08 | *n.s* | 0.53±0.05 | 0.97 |

*N.D*, not detectable, *Adamts*, A Disintegrin And Metalloproteinase with Thrombospondin Motifs; *Ctsd*, Cathepsin D; *Ccbe1*, collagen- and calcium-binding EGF-like domains 1; *Pcsk*, proprotein convertase; *Flt4*, VEGFR3. Data reported as fpkm (mean ± sem). FC, fold of control. DESeq2 Wald test comparing control to Il1β-treated LECs. For full list of DEGs, see supplementary **Table S5**.

#### Suppl. Table 4 – Cardiac gene expression at 8 weeks post-TAC

| gene | Sham N=6 | Sham p | TAC N=8 | TAC + *early* N=7 | TAC + *early* p | TAC + *late* N=7 | TAC + *late* p |
|---|---|---|---|---|---|---|---|
| ***Lyve1*** | **1.00**±0.05 | *n.s* | **1.14**±0.18 | **1.41**±0.17 | *n.s* | **2.20**±0.17 | *** |
| ***Pdpn*** | **1.00**±0.11 | *** | **2.21**±0.28 | **2.25**±0.35 | *n.s* | **2.12**±0.09 | *n.s* |
| ***Ccl21*** | **1.00**±0.09 | *n.s* | **1.21**±0.14 | **1.44**±0.23 | *n.s* | **1.91**±0.13 | * |
| ***Vegfc*** | **1.00**±0.06 | * | **1.65**±0.13 | **1.82**±0.18 | *n.s* | **1.71**±0.08 | *n.s* |
| ***Vegfd*** | **1.00**±0.09 | *n.s* | **1.48**±0.3 | **1.12**±0.08 | *n.s* | **0.88**±0.08 | * |

Data reported as fold of sham (mean ± sem). All samples were compared using Two-way ANOVA followed by Sidak's posthoc test. * p<0.05; ** p<0,001; *** p<0.001 *versus* TAC controls.

### Suppl. Methods & Materials

#### Human samples

Five micrometer thick sections were prepared from paraffin-embedded human septal samples. Standard immunohistological protocols, including citrate antigen retrieval, were used to reveal podoplanin-positive lymphatic vessels using a mouse anti-human podoplanin/D2-40 antibody validated for clinical practice (Dako, # M3619, diluted *1:50*). Signal was revealed following incubation with HRP-conjugated donkey anti-mouse secondary, followed by DAB detection kit (Vector Laboratories, Burlingame, CA). All patient slides were processed together for uniformity. Lymphatics were examined using a Zeiss axiovision light microscope equipped with CCD cameras at *x10.* Lymphatic vessels, identified as Podoplanin$^+$ vascular structures in the subendocardium, were counted by an observer blinded to the patient category. Total lymphatic density, open lymphatic density (lumenized vessels), lymphatic vessel diameter, and density of perivascular lymphatics (defined as lymphatics within 200 µm distance of a large blood vessel) were determined. On average for each patient 1.2 ±0.2 mm² of the septal sample was analyzed to determine lymphatic vessel sizes and densities.

#### Experimental Model

Female BALB/c mice (22-24 g) were obtained from Janvier. Animal housing and experiments were in accordance with European Directive 2010/63/EU on the protection of animals, and the study was approved by the Normandy University ethical review board Cenomexa according to French and EU legislation (APAFIS #**23175**-2019112214599474 v6; APAFIS #**32433**-2022070712508369 v2). Minimally-invasive transversal aortic banding constriction (TAC) was performed on 8 week-old mice, as previously described.[1] Mice were anaesthetized by intraperitoneal injection of ketamine (100 mg kg $^{-1}$ Imalgene®) and xylazine (10 mg kg $^{-1}$ Rompun® 2%, Bayer Health Care) and placed on mechanical ventilation. The operator performed a minimal thoracotomy with an incision at the level of the first intercostal space. The aortic arch was visualized under low-power magnification. A snare, made of 7-0 polypropylene suture, was passed under the aorta between the origin of the right innominate and left common carotid arteries. Two suture bands were placed side-by-side to create an elongated stenosis and prevent internalization of the suture, as described.[2] A bent 26-gauge needle was placed next to the aortic arch, and the sutures were snugly tied around the needle and the aorta. After banding, the needle was quickly removed. The skin was closed, and mice recovered on a warming pad until fully awake. The sham procedure was identical except that the aorta was not banded. Buprenorphine (50 µg/kg, Buprecare®, Axcience) was injected subcutaneously 6 hours after surgery and twice per day until 3 days post-operation. Anti-IL-1β treatment consisted of a blocking monoclonal antibody against IL-1β (Gevokizumab/XOMA-052 provided by Servier, France) as previously described.[3] The antibody was administered by repeated intraperitoneal injections of 200 µg/mouse (10 mg/kg$^{-1}$) three times per week starting from week-1 until week-5 post-TAC (*early* treatment) or starting from week-4 until week-8 post-TAC (*late* treatment). Euthanasia was performed by pentobarbital overdose (100 mg kg$^{-1}$ Euthoxin).

#### *Echocardiography*

Transthoracic echocardiography measurements (Doppler and M-mode) were performed in animals anaesthetized with Isoflurane (1-2%), using a Vevo 3100 Imaging System (FUJIFILM). The heart was analysed in the two-dimensional mode in parasternal short-axis views. LV fractional shortening and ejection fraction were calculated using M mode imaging. To evaluate heart rate (HR) and velocity time integral (VTI), a pulsed Doppler of the LV outflow tract was performed, allowing calculation of stroke volume ($SV = \pi \times$ LV outflow radius$^2 \times VTI$) and cardiac output ($CO = SV \times HR$). LV outflow radius was measured for each animal. Aortic flow was measured using pulsed Doppler to verify the efficacy of transversal aortic constriction compared to sham mice. Echocardiographic analyses were performed with VevoLAB software.

##### *Immunohistochemistry*

Murine cardiac samples were sectioned into a central slice, which was snap-frozen. Cardiac sections were cut on a cryostat (8 µm thickness) and collected on SuperFrost plus glass slides. After fixation in acetone for 10 min, non-specific binding sites were blocked in 3% BSA in PBS, followed by Biotin-Avidin Blocking kit (Thermo Scientific) when streptavidin (SA)-conjugates were used to detect biotinylated secondary antibodies. Primary antibodies (*see Suppl. Table 3*), diluted in 1% BSA in PBS, were incubated on the sections at r.t. for 1h, followed by repeated washing in PBS and incubation with secondary antibodies (*see Suppl. Table 5*) for 30 minutes to 1h. Multi-stainings were performed sequentially, and negative controls included omission of primary antibodies. Slides were mounted in Vectashield containing DAPI, and images were acquired using x20 or x40 objectives on a Zeiss epifluorescence microscope (AxioImager J1) equipped with an apotome and Zen 2012 software (Zeiss). Images were analyzed by an operator blinded to the treatment groups using Fiji imaging software (NIH).

##### Suppl. Table 5 – antibodies used for immunohistochemistry in mice

| antigen | article | supplier | species reactivity | host | dilution | Conc (µg/mL) |
|---|---|---|---|---|---|---|
| **CCL21** | *AF457* | RnD Systems | *mouse* | **goat** | 1/100 | **5** |
| **CD68** | *14-0681-82* | eBioscience | *mouse* | **rat** | 1/800 | **5** |
| **CD206** | *PA5-46994* | ThermoFisher Scientific | *mouse* | **goat** | 1/800 | **0.25** |
| **CCR2** | *273050* | Abcam | *mouse* | **rabbit** | 1/600 | **0.75** |
| **IL-1β** | *AF401NA* | RnD Systems | *mouse* | **goat** | 1/100 | **2** |
| **LYVE-1** | *103-PA50* | Reliatech | *mouse* | **rabbit** | 1/1000 | **0.4** |
| **Biotinylated LYVE-1** | *103-PA50-bi* | Reliatech | *mouse* | **rabbit** | 1/500 | **0.8** |
| **Podoplanin** | *14-5381-82* | eBioscience | *mouse* | **hamster** | 1/10 000 | **1** |
| **WGA-FITC** | *FP-CE8070* | Interchim | | | 1/100 | **1** |
| **antigen** | **article** | **supplier** | | **Fluorochrome** | | **Conc (µg/ mL)** |
| **Donkey anti-Rabbit** | *711-165-152* | Jackson Immunoresearch | | **Cy3** | | **3** |
| **Donkey anti-Rabbit** | *711-606-152* | Jackson Immunoresearch | | **AF647** | | **3** |
| **Streptavidin** | *FP-CA5640* | Interchim | | **Fluoprobe 647** | | **0.7** |
| **Streptavidin** | *FP-XEQ270* | Interchim | | **Cy7** | | **0.7** |
| **Donkey anti-Rat** | *712-546-150* | Jackson Immunoresearch | | **AF480** | | **3** |
| **Donkey anti-Rat** | *712-586-152* | Jackson Immunoresearch | | **Cy3** | | **3** |
| **Donkey anti-Goat** | *A50-201D3* | Interchim | | ***DYLIGHT 550*** | | **1.25** |
| **Biotinylated Donkey anti-Goat** | *705-605-147* | Jackson Immunoresearch | | | | **1.9** |
| **Goat anti-Hamster** | *A-21110* | ThermoFisher Scientific Invitrogen | | **AF488** | | **0.8** |

##### *Image analysis of cardiac sections*

Murine lymphatic vessels were defined as strongly Lyve1-positive structures, lacking macrophage marker (CD68), and mostly also positive for Podoplanin. Lymphatics are preferentially located in the subepicardium in rodent hearts. These Lyve1-positive, CD68-negative vessels clearly differ from $CD68^+$ macrophages, which either lack or display weaker Lyve1 expression. These macrophages, also differing in size and morphology from the elongated cell body and nucleus typical of lymphatic endothelial cells, were readily excluded from lymphatic counts. Photos were captured at *x20*.

Due to the absence of conclusive markers to distinguish precollectors from capillaries in murine cardiac lymphatics, we made use of the fact that precollectors preferentially run in a base-to-apex fashion, whereas lymphatic capillaries lack such consistent spatial organization. Hence, by evaluating vessels with a lumen parallel to coronal cardiac sections, the precollector population of vessels is more frequently detected as vessels running perpendicular to the section (= open lumen vessels). To assess the size and frequency of open lymphatic vessels (diameter > 5 µm; a population enriched in precollector vessels), between 10-20 images were captured at *x20* from the LV free-wall were analyzed for each mouse. The density ("open vessels/mm²") and lymphatic lumen sizes were measured and used to calculate mean vessel diameter. The parameter "% open lymphatic area" was calculated as the sum total of all measured lymphatic diameters divided by the total cardiac area analyzed for each mouse heart. On average for each mouse 2.2±0.3 mm² of the LV was analyzed to determine lymphatic vessel sizes and % open lymphatic area.

To assess perilymphatic chemokine gradients, we used ImageJ to outline lymphatic lumen, and then subtract intralymphatic Ccl21 signal from the image (see **Suppl. Fig. 6a**), and assess the area surrounding each lymphatic vessel for area % of Ccl21 positivity (set as a minimum of 15 levels of grey).

Cardiomyocyte sizes were evaluated in sections double stained for CD31 and WGA imaged at *x40*.

Cardiac macrophages were analyzed in images captured at *x20*. Total cardiac macrophages were defined as CD68 expressing cells. *Cardiac-resident macrophages* were defined as $LYVE1^{+}$ cells, whereas infiltrating macrophages were defined as $LYVE1^{neg}$ cells. Among these, cardiac-recruited $CCR2^{+}$ $CD206^{neg}$ $LYVE1^{neg}$ macrophages were denoted "*classical infiltrating macrophages"*, whereas $CD206^{+}$ $LYVE1^{neg}$ macrophages were denoted "*alternative infiltrating macrophages*".

##### *Histology in mouse or human cardiac sections*

Cardiac cryosections (8 µm) of mouse samples, or paraffin sections (5 µm) of human samples, were processed for Sirius Red staining, as described.[1]

Cardiac interstitial collagen density was evaluated in mouse cardiac sections imaged on a light microscope (Zeiss) equipped with an *x40* objective.

Replacement fibrosis (micro-scars) was evaluated in these images as total number/cardiac section of fibrotic hotspots, identified as images containing >11% fibrotic area per image, analyzing on average 14±0.5 images/sample (average total examined area 0.5 mm² of each LV cross-section).

Perivascular fibrosis was evaluated in images taken at *x20,* focused on arterioles with a lumen diameter of 16-60 µm in both mice and humans. On average 5 vessels/cardiac section were used to determine perivascular fibrotic area (µm²), as previously described.[4] Images were analyzed by an operator blinded to treatment groups using Fiji imaging software (NIH).

Briefly, for each arteriole, three areas were measured using ImageJ software: the luminal area, the total arteriolar area, and the total fibrotic area (see **Suppl. Fig. 6b**). The perivascular fibrosis area was calculated by subtracting the total arteriole area from the total fibrotic area. The luminal area was also used to determine arteriole diameter, allowing classification of arterioles by size.

##### *Real-Time Polymerase Chain Reaction*

Murine cardiac samples were collected at 8-weeks post-TAC. RNA extraction was performed with Trizol method. Briefly, samples were homogenized (45 sec.) in TRIZOL using Precellys tubes with beads. RNA was obtained after incubation, successively, in chloroform and isopropanol and three washes in ethanol. RNA quality was analyzed by Nanodrop 2000 (Thermo Fisher Scientific). cDNAs were generated using a reverse transcriptase after DNAse treatment. Real-time PCR was performed on a LightCycler 480 (Roche Molecular Biochemicals) with a commercially available mix (FastStart DNA Master SYBR Green I kit; Roche) on white 96-well plates. The reactions were performed in duplicate for each sample. Concentrations were calculated using a standard curve, obtained by serial dilutions of samples from an organ with high expression of the gene of interest. To normalize gene expression, values are expressed as ratios of a reference gene (E2F). Healthy sham animal ratios are then set as 100%, and values in TAC-operated animals are expressed relative to their respective sham controls. The sequences of the specific primers are detailed in *Suppl. Table 6.*

##### Suppl. Table 6 – list of primers used in mouse cardiac samples for qPCR

| gene | sense | antisense | amplicon (bp) |
|---|---|---|---|
| ***Eef2*** | GCGAGGACAAAGACAAGGAG | GGGATGGTAAGTGGATGGTG | 108 |
| ***Ccl2*** | CCCAATGAGTAGGCTGGAGA | GCTGAAGACCTTAGGGCAGA | 210 |
| ***Ccl7*** | CAGAAGGATCACCAGTAGTCGG | ATAGCCTCCTCGACCCACTTCT | 108 |
| ***Ccl21*** | TCCCTACAGTATTGTCCGAGGC | ATCAGGTTCTGCACCCAGCCTT | 141 |
| ***Cxcl12*** | GGAGGATAGATGTGCTCTGGAAC | AGTGAGGATGGAGACCGTGGTG | 133 |
| ***Col1a*** | CCTCAGGGTATTGCTGGACAAC | CAGAAGGACCTTGTTTGCCAGG | 115 |
| ***Flt4*** | AGACTGGAAGGAGGTGACCACT | CTGACACATTGGCATCCTGGATC | 128 |
| ***Il1β*** | TGGACCTTCCAGGATGAGGACA | GTTCATCTCGGAGCCTGTAGTG | 148 |
| ***Il6*** | TACCACTTCACAAGTCGGAGGC | CTGCAAGTGCATCATCGTTGTTC | 116 |
| ***Lyve1*** | ACCAGGTAGAGTCAGCGCAGAA | CAGGACACCTTTGCCATTCTTCC | 128 |
| ***Nppb*** | TCCTAGCCAGTCTCCAGAGCAA | GGTCCTTCAAGAGCTGTCTCTG | 98 |
| ***Pdpn*** | ACAACCACAGGTGCTACTGGAG | GTTGCTGAGGTGGACAGTTCCT | 116 |
| ***Tgfb1*** | TGATACGCCTGAGTGGCTGTCT | CACAAGAGCAGTGAGCGCTGAA | 107 |
| ***TNFα*** | TAGCCAGGAGGGAGAACAGA | TTTTCTGGAGGGAGATGTGG | 139 |
| ***Vegfc*** | AGCCCACCCTCAATACCAG | GCTGCTCCAAACTCCTTCC | 154 |
| ***Vegfd*** | CTCCACCAGATTTGCGGCAACT | ACTGGCGACTTCTACGCATGTC | 112 |

##### *Western blot analysis in cardiac samples*

Cardiac protein was extracted using Cell lysis buffer (#895347, RnD Systems) in microbead tissue dissociation tubes (Precellys #P000933-LYSK0-A – 0.5 mL). Twenty-five micrograms of denatured cardiac protein from healthy sham-operated or TAC-operated mice at 8 weeks post-TAC treated or not with anti-IL-1β antibodies, were separated on stain-free 4-20% agarose gels (Criterion TGX Precast Gel, Biorad), followed by blotting with a rabbit anti-VEGFC antibody (#ab9546, Abcam; *1:1000*), or a rabbit anti-VEGFD antibody (#ab155288, RnD systems; *1:500*), and finally a mouse anti-actin antibody (#15597191, Invitrogen; *1:10000*) diluted in PBS/BSA 5%. After repeated washing, HRP-conjugated secondary antibodies (goat anti-rabbit #P0448, Dako; *1:5000*, donkey anti-mouse #A16011, Invitrogen; *1:5000*) were used. Target proteins were visualized using ECL chemiluminescence kit (#1705061, BioRad). Ratios of specific bands to actin signal per well was analyzed in ImageJ. For VEGF-C and VEGF-D multiple bands were present, representing the various cleaved forms of the growth factors. For VEGF-C, the ratio of mature to immature forms was calculated as the ratio of the densities of cleaved, mature monomeric VEGF-C (21 kDa bands) to the densities of full-length immature monomeric pro-VEGF-C (31 kDa bands)[5]. For VEGF-D only the full-length immature pro-VEGF-D form (42 kDa dimer) was visible (cleaved, mature form at 32 kDa too weak for analysis). VEGF-C and VEGF-D protein ratios were compared between sham and TAC groups using two-way ANOVA followed by Sidak's posthoc test.

Samples of macrophage-conditioned media (average protein concentration 1 mg/mL) were prepared with β-mercapto, and 23 micrograms of denatured protein from control or IL-1β-treated cells (*n=6* samples/group) were separated on 4-15% precast polyacrylamide gels (Mini-PROTEAN TGX Gels, BioRad), followed by blotting with a rabbit anti-VEGFC antibody (#ab9546, Abcam; 1:200) diluted in TBS-T/BSA 5%. After repeated washing, biotin-conjugated secondary antibodies (biotinylated goat anti-rabbit; 111-065-144, Jackson ImmunoResearch, 1:5000) was used, followed by SA-HRP (N100, Thermoscientific, 1:5000). Subsequently, blots were incubated with a mouse anti-actin antibody (MA5-15739, Invitrogen; 1:5000), followed by a secondary antibody (HRP-conjugated donkey anti-mouse #A16011, Invitrogen; 1:10000). Target proteins were visualized using ECL chemiluminescence kit (#170-5061, BioRad). Total protein content per lane was evaluated by colorimetric staining for Ponceau red. Ratios of specific bands of VEGF-C to actin signal per lane were analysed in ImageLab. For VEGF-C, multiple bands were present: a full-length (immature) 31 kDa band and 15-21 kDa cleaved (mature) monomeric band. Levels of each protein form, and the relative ratio of mature to immature VEGF-C, were calculated and presented as fold of control. Data were compared between control and IL-1β treated groups using two-way ANOVA followed by Sidak's posthoc test.

##### *Macrophage cell culture*

Bone marrow-derived macrophages (BMM) were prepared from femur and tibia of C57/BL6 mice. Briefly, bone marrow-derived cells were seeded in 6-well plates ($6\times10^6$/well) and differentiated into macrophages in RPMI1640 medium containing 15% L929-conditioned medium, 10% fetal calf serum (*FCS*, heat inactivated at 56 °C/ 45min), 50 µM β-mercaptoethanol, 100 U/mL penicillin, and 100 µg/mL streptomycin during 7 days, as described[6]. On day 7, BMMs were serum-starved in RPMI medium containing 0.5% FCS for 4h and exposed for 4 and 24 hours to 10 ng/mL recombinant murine IL-1β (PreproTech, 211-11B) in 0.6 mL per well of 0.5% FCS cell culture media. Total RNA was isolated using NucleoSpin RNA kit (Macherey Nagel, 740955) according the instruction. RNA was transcribed using First Strand cDNA Synthesis Kit (Thermo Scientific, K1612). qPCR was done on GeneQuant Studio 6 Real-Time PCR System using SybrGreen PCR Mix. Primers used in qPCR are listed in *Suppl Table 7*.

##### Suppl. Table 7 – list of primers used in mouse macrophages for qPCR analysis

| gene | sense | antisense | amplicon (bp) |
|---|---|---|---|
| ***Adamts3*** | TTCCAGGAACCTCTGTTGCC | GCTGATCTCTTGTAGACAAC | 151 |
| ***Adamts14*** | CTGATCATGGTGGGCTACCGACA | CCATCCTCGTGGTTGAGGGCACA | 236 |
| ***Ctsd/CatD*** | AGGTGAAG GAGCTGCAGAAG | ATTCCCATGAAGCCACTCAG | 199 |
| ***Ccbe1*** | ATGGGACCTATGGGACCTTC | AGTGAGTCCGGTGTCCAAAC | 235 |
| ***Furin*** | TGAGCCATTCGTATGGCTACG | GGACACAGCTTTTCTGGTGCA | 576 |
| ***Pcsk5*** | AGGTGGAGTGGATCCAACAG | CCGTGTAGCCTCTCTTCCAG | 189 |
| ***Pcsk7*** | CCACCCTGATGAGGAGAATG | GCCACAGCCTCCATACTGTC | 165 |
| ***Vegfc*** | CGAGGTCAAGGCTTTTGAAG | TCCCCTGTCCTGGTATTGAG | 165 |

*Adamts*, A Disintegrin And Metalloproteinase with Thrombospondin Motifs; *Ctsd*, Cathepsin D; *Ccbe1*, collagen- and calcium-binding EGF-like domains 1; *Pcsk*, proprotein convertase

##### *Lymphatic cell culture*

Balb/c dermal LECs (CellBiologics, BALB-5064L) were seeded in 6-well plates ($3\times10^5$/well) and grown to confluence in complete Endothelial Cell Medium (ECM, CellBiologics, M1168) including 5% FCS with VEGF, ECGF, EGF supplement, according to recommendations of the supplier. Next, cells were serum-starved (1% FCS, ECM w/o VEGF, ECGF, EGF) overnight, and exposed to 20 ng/mL recombinant mouse IL-1β (PeproTech) in 2 mL per well of serum-starvation media. After 24h cells were recovered and RNA extracted (RNeasy minikit, 74104, Qiagen) for subsequent mRNAseq analyses and bioinformatics analyses (X204SC23121183-Z02-F002) by NOVOGENE company (Cambridge, UK) for identification of differentially expressed genes. Briefly, poly-A mRNA-seq, sequencing length of 150 nt paired-end (PE150), were performed with Illumina Seq. Reads were aligned to the mouse genome (GRCm38). Transcript quantification was performed with featureCounts, and differential analysis, including normalization of the original *readcount*, was performed with DESeq2 bioconductor package. Statistical test (DESeq2 Wald test) was conducted on the expression matrix after standardization. Genes significantly up- or downregulated upon treatment were identified (for RNAseq data on lymphangiogenesis regulators see *Suppl Table 3*), and results are reported as Fpkm and as fold of control condition (1% FCS in ECM without cytokines and growth factors). The original data included in this study has been deposited in GEO (GSE290146, *Effect of interleukin-1beta on confluent murine lymphatic endothelial cells*).

#### Suppl. References
